# ReSeT: a taxonomy-aware reference genome selection tool

**DOI:** 10.64898/2026.06.17.732946

**Authors:** Jasper van Bemmelen, Jasmijn A. Baaijens

**Affiliations:** Intelligent Systems Department, Delft University of Technology, Van Mourik Broekmanweg 6, 2628 XE, Zuid-Holland, Netherlands

**Keywords:** taxonomic profiling, facility-location problem, local search, genome selection

## Abstract

**Motivation:** Reference genome composition determines which taxa a profiling pipeline can detect and distinguish, and becomes of critical importance for high-resolution profiling where taxonomic boundaries begin to blur. Existing selection tools optimize within-taxon representativeness but disregard discrimination across taxa, leaving open whether explicitly accounting for inter-taxon discrimination during selection improves profiling.

**Results:** Here we present ReSeT, a facility-location-based reference genome selection tool that operates on arbitrary pairwise distance matrices, extended with a tunable inter-taxon discrimination term and per-genome selection cost, and solved by local search. We benchmark ReSeT against established selection methods on three viral datasets spanning varying degrees of taxonomic ambiguity. On the high-ambiguity SARS-CoV-2 datasets, appropriately tuned ReSeT selections matched or exceeded the strongest alternatives in terms of profiling accuracy, whereas on the low ambiguity IAV dataset VSEARCH remained dominant. Interestingly, we find that the novel inter-taxon discrimination term contributed weakly, indicating that ReSeT’s facility-location formulation and selection cost drives ReSeT’s performance. We further propose a novel taxonomic ambiguity index, computable from ReSeT’s inputs, that summarizes the taxonomic ambiguity of reference genomes and aligns with where ReSeT improves over existing selection methods.

Availability and implementation: ReSeT is implemented in Python (≥3.10) and is freely available under the MIT license. The source code is available on GitHub at https://github.com/JaspervB-tud/ReSeT and ReSeT can also be installed directly from the Python Package Index (PyPI) via pip install reset-bio.

## Introduction

Reference genomes play a central role in taxonomic profiling pipelines: they define which taxa can be detected, and how well those can be distinguished from others. When reference taxa are well-separated, which is often the case at lower taxonomic resolutions (i.e. species-level and above), this dependence is of little consequence. In this case, including complete taxonomic databases, such as RefSeq [1], may suffice as the primary objective is to cover diversity within each taxon. But with rising interest in high-resolution profiling (e.g. bacterial strain-level), motivated by the functional and phenotypic variability observed between strains within species [2], taxon boundaries begin to fade [3, 4] and discrimination between taxa becomes increasingly difficult. Under these conditions, explicit reference genome selection provides a way of dealing with the reduced taxonomic separation: in previous work we have shown that generic sequence dereplication tools can be applied to the database to significantly improve accuracy for both bacterial strain-level (*Escherichia coli*) and viral lineage-level (SARS-CoV-2) profiling [5].

In practice, reference genome selection relies on a range of tools that vary considerably in design. General-purpose dereplication methods such as VSEARCH [6], MMseqs2 [7] and MeShClust [8] are routinely repurposed for this task, using their clustering-based representatives as candidate references. dRep [9] and GGRaSP [10] were specifically developed for genome selection and perform hierarchical clustering, but can further incorporate assembly quality criteria to obtain high-quality references. More closely related to our work, PARNAS [11] and Repset [12] formulate selection as facility-location and submodular optimization problems, but are restricted in their applicability: PARNAS requires genetic distances to form a tree metric, limiting application to cases where a phylogenetic tree is available, and Repset relies on (PSI-)BLAST [13] for distance computations, which becomes prohibitive at genome scale. Despite differences in their design and implementation, these methods all share a structural limitation: they only optimize representativeness within groups of sequences, but do not consider discrimination across taxa, leaving open whether explicitly incorporating inter-taxon discrimination can further improve profiling, and under what conditions.

Here we introduce ReSeT (**Re**ference genome **Se**lection **T**ool), a reference genome selection method formulated as a facility-location problem (FLP) on general pre-computed pairwise distance matrices, extended with a tunable inter-taxon discrimination term and per-genome selection cost. The resulting objective is optimized via local search, and by operating on general distance matrices, ReSeT removes the input constraints of existing FLP-based approaches, allowing direct incorporation of standard sequence (dis)similarity estimation tools like MASH [14] and sourmash [15]. The tunable inter-taxon term allows us to directly test how explicitly penalizing similarity across taxa during selection affects downstream profiling accuracy.

We benchmark ReSeT against existing selection strategies on two viruses with distinct levels of taxonomic ambiguity: SARS-CoV-2 (weakly separated), and influenza A virus (strongly separated). On SARS-CoV-2, appropriately tuned ReSeT configurations outperform the strongest alternatives, while on IAV no method consistently dominates across metrics and difficulties. Analysis of ReSeT’s objective components shows that intra-taxon coverage and selection cost strongly drive performance, while the inter-taxon discrimination term contributes weakly – even in the high-ambiguity regime where it would be expected to help most. This suggests that the FLP formulation itself, rather than inter-taxon penalization, brings the main improvement over other approaches. Our second contribution is an ambiguity index, directly computable from ReSeT’s input, that quantifies how many genomes are closer to taxa other than their own in a given reference database. In our datasets, ReSeT’s advantage was observed when this index was large (i.e. taxa weren’t well-separated), while VSEARCH was superior when the index was small. These results suggest that it can serve as a pre-screening criterion for ReSeT-based selection.

## Methods

### ReSeT

ReSeT finds a set of reference genomes by solving a variant of the facility location problem (FLP) [16], a combinatorial optimization problem in which one selects a subset of *facilities* from a set of candidates to serve a set of *clients*, minimizing the total distance from each client to its nearest selected facility, plus a fixed cost per opened facility. In our setting, every candidate reference genome is both a facility and a client, and distances are computed from sequence similarity. Unlike existing FLP-based sequence selection tools [11, 12], ReSeT operates on general pairwise distance matrices on a [0, 1] range, and extends the standard FLP objective with a per-taxon feasibility constraint and a tuneable inter-taxon separation term that explicitly penalizes selections obfuscating taxon boundaries.

#### Notation

Let *G* denote the set of candidate genome assemblies, *T* the set of taxa, and *τ* : *G* → *T* a taxon assignment function. We assume a pairwise distance function *d* : *G × G* → [0, 1] that is symmetric (*d*(*g, g*^*′*^) = *d*(*g*^*′*^, *g*)) and reflexive (*d*(*g, g*) = 0 ∀*g* ∈ *G*), but is not required to satisfy any additional properties, allowing compatibility with most practical sequence distance estimators. The [0, 1] range aligns with common genetic distance formulations, which are usually defined as dissimilarities, and supports the use of 1 − *d*(*g, g*^*′*^) as a similarity in the inter-taxon penalty below. A feasible selection is a subset *G* ⊆ *G* containing at least one genome from each taxon *t* ∈ *T*. For convenience, we let *G*_*t*_ := *{g* ∈ *G* : *τ* (*g*) = *t}* denote the set of genomes in *G* ⊆ *G* that are from taxon *t* ∈ *T*, and assume an arbitrary total order on *T*, so that *{t* ∈ *T* : *t < t*^*′*^*}* denotes all taxa preceding *t*^*′*^.

#### Intra-taxon coverage

Given a feasible selection *G* ⊆ *G*, we incur a penalty for each unselected genome *g* ∈ *G \ G* equal to its distance to the closest selected genome of the same taxon:

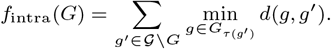

This promotes the selection of genomes that maximize similarity to unselected genomes on a per-taxon basis.

#### Inter-taxon separation

To discourage selecting representatives that obfuscate taxonomic boundaries, we impose an inter-taxon penalty for all distinct taxon pairs. Specifically, for each distinct pair (*t, t*^*′*^), we penalize the most similar (least distinct) selected inter-taxon pair as follows:

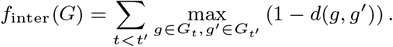

This term, along with *f*_intra_ implicitly creates a trade-off between selecting genomes to cover taxon heterogeneity and preserving taxon separation. Moreover, the use of the maximum operator (rather than e.g. the mean of inter-taxon similarities) preserves linearity, allowing the problem to be represented as a mixed-integer linear program (see Supplementary Material).

#### Complete objective function

In addition to *f*_intra_ and *f*_inter_, we also impose a selection cost *c* which assigns a fixed penalty per selected genome, to control the size of the resulting reference set. Larger *c* yields smaller, more compact selections, while smaller *c* allows more references at the cost of increased redundancy. The meaningful range of *c* depends on the scale of the chosen distance function *d*: if distances cluster near 0 (i.e. all genomes are highly similar), even small values of *c* can trivially force the selection to at most one genome per taxon. Practical use of *c* therefore requires choosing it on the same scale as typical distances under the chosen distance function *d*.

Combining *f*_intra_, *f*_inter_ and the selection cost *c* into a single objective requires accounting for a structural imbalance between these terms: in many practical settings there is a substantial difference between the number of taxa |*T* | and the number of available genomes |*G*|, leading to a skewed weighting of *f*_intra_ and *f*_inter_ when combined as-is. To correct for this, we instead consider the following weighted objective function:

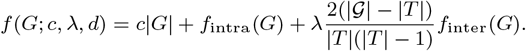

Here, the inter-taxon component is first rescaled by 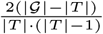 to account for the difference in intra- and inter-taxon terms in the objective. Afterwards, the rescaled inter-taxon term is multiplied by a user-defined scalar *λ* to control the impact of the inter-taxon term in the objective.

##### Local search implementation

The problem defined above is NP-hard in general (see Supplementary Material), making exact optimization infeasible in practice. Both prior FLP-based selection tools sidestep this through restrictive assumptions on their inputs and objectives: PARNAS achieves polynomial-time optimality by requiring *d* to be a tree metric, and Repset utilizes a greedy approximation algorithm by exploiting the submodular nature of the FLP-based selection problem without inter-taxon penalization. ReSeT’s combination of general distance matrices with a non-submodular objective breaks both these assumptions, and we instead optimize the objective using a local-search heuristic that iteratively applies objective-improving moves to find a locally optimal solution.

#### Initialization

We initialize a feasible solution by selecting a random fraction of genomes, forcing at least one representative per taxon.

#### Neighborhood definition

Given a feasible solution *G*, we define its neighborhood *N* (*G*) as the set of feasible solutions that can be obtained from *G* by applying one of the following moves:

1. **Add:** add an unselected genome *g* ∈ *G \ G*,
2. **Remove:** remove a selected genome *g* ∈ *G*, provided that |*G*_*τ*(*g*)_| *>* 1,
3. **1-Swap:** within a taxon *t*, add an unselected genome and remove one selected genome,
4. **2-Swap:** within a taxon *t*, add two unselected genomes and remove one selected genome.

These operations are common in local search algorithms for facility location problems and, in principle, allow transitions between any pair of feasible solutions [17]. Although arbitrary swap sizes could further increase the local search’s flexibility, adding them leads to exponentially large neighborhoods (Supplementary Material), while restricting to 1- and 2-swaps instead maintains some of the flexibility but keeps neighborhood evaluations computationally tractable.

#### Move evaluation and termination criteria

For optimization, we use a *first-improvement local search with randomized move order* where, in every iteration, we evaluate moves in the current neighborhood in randomized order, accepting the first improving move found. All move types defined here maintain feasibility, thus we do not need to impose additional penalties for moving to infeasible solutions. The algorithm terminates when (i) no improving moves can be found (i.e. the solution is locally optimal), (ii) a maximum number of iterations is reached, or (iii) the maximum allowed wall-clock runtime is exceeded. After termination, the best solution found is returned.

To speed up the local search routine, we also incorporate a user-defined threshold that defines the amount of time the algorithm may spend on an iteration before removing 2-swaps from the move pool. When this threshold is exceeded, the algorithm will only allow **add, remove** and 1**-swap** moves for the remainder of the iteration, sacrificing strict local optimality for computational efficiency.

### Experimental design

We evaluate ReSeT against established reference selection methods on three viral datasets varying in taxonomic ambiguity: two SARS-CoV-2 datasets (USA- and China-based) with high ambiguity, and one influenza A virus (IAV) dataset with low ambiguity. We compare against four baselines (applied per taxon) which were shown to perform well in prior work [5]: VSEARCH v2.28.1 [6], which performs greedy incremental clustering; hierarchical clustering with complete-linkage (SciPy v1.5.0, https://scipy.org) selecting medoids per cluster as representatives; medoid selection, retaining only the medoid genome per taxon; and a no-selection baseline retaining all reference genomes. Despite their similarity to ReSeT, we omitted PARNAS due to requiring a phylogenetic tree, and Repset [12] because its runtime on the single largest SARS-CoV-2 lineage exceeded 1.5 days.

#### Datasets

Our SARS-CoV-2 datasets consist of geographically restricted genomes from either the USA or China, downloaded from GISAID [18] on November 16th, 2025. For both datasets, genomes had a collection date between January 1st 2022 and June 30th 2024, and genomes were filtered based on ambiguous nucleotides, length, and host. To maintain computational feasibility, we further downsampled the USA dataset resulting in 85,760 genomes over 1,474 Pango lineages [19] in the USA dataset, and 11,862 genomes over 356 lineages in the China dataset. Filtering and downsampling details can be found in the Supplementary Material.

We also consider an IAV dataset consisting of all available high-quality HA and NA segments of the H1N1pdm09 and H3N2 subtypes, with collection dates between January 1st 2024 and March 31st 2025, downloaded from GISAID [18] on November 20th, 2025. We applied the same filtering rules to these sequences (except for length filters), including downsampling, resulting in a dataset of 29,909 genomes (represented by only the constituent HA and NA segments) over 8 clades.

In our experiments, we partition the data into a reference pool with candidate reference genomes, and a simulation pool with candidate genomes for read simulations based on a temporal cut-off (see Supplementary Material). This split ensures that there is no leakage of simulation genomes into reference sets, and mimics reality where reference genomes are obtained before new samples are analyzed.

#### Sample design

We evaluate profiling accuracy under two complementary sample regimes: difficulty-stratified samples that isolate the effect of taxonomic ambiguity, and Dirichlet-distributed samples representing randomized compositions.

##### Difficulty-stratified samples

To isolate the effect of taxonomic ambiguity on profiling performance, we construct samples at three difficulty levels (Easy, Medium, Hard) for SARS-CoV-2, and two (Easy, Hard) for IAV. We treat a genome as a source of ambiguity when it is more similar to genomes of another taxon than to those of its own, and we stratify difficulty levels within each dataset by how much of this ambiguity they contain: Easy samples minimize it, Hard samples maximize it, and Medium samples (when defined) sit in between. Due to differences in taxonomic structure across the two viruses, we operationalize this stratification differently for each, as described below.

For SARS-CoV-2, we align all simulation genomes with MAFFT v7.525 [20] to compute pairwise genomic distances from the resulting MSA, then cluster genomes with HDBSCAN v0.8.39 [21] (see Supplementary Materials for details). Within each cluster, we compute per-lineage medoid genomes, and label a genome *difficult* if its nearest lineage medoid is from another lineage. However, if two lineages *T* and *T*^*′*^ are both present in a sample, and each contains a genome closer to a medoid of the other, a read from *T* misassigned to *T*^*′*^ can be compensated by misassigning a read from *T*^*′*^ to *T*, resulting in no observable error. We therefore build an undirected *lineage difficulty graph* with an edge between two lineages whenever a difficult genome in one has its nearest medoid in the other within at least one cluster. From the graph, we select up to 10 lineages from a maximum-weight independent set ensuring that no two selected lineages will mutually absorb each other’s misclassifications. From the selected lineages, Hard samples retain up to 5 difficult genomes per lineage. Medium samples also retain up to 5 genomes per selected lineage, but prioritize non-difficult genomes instead. Finally, Easy samples retain up to 5 genomes per lineage from up to 10 lineages isolated in the difficulty graph.

For IAV, with only 8 clades and limited inter-clade ambiguity, the graph-based approach above is unstable: depending on HDBSCAN parameters, clustering labels either every clade or no clade as containing difficult genomes. We therefore stratify at the genome level rather than the clade level, retaining all clades. We compute per-clade medoid genomes globally, and score each genome by the difference between its distance to its own clade medoid and to the nearest other-clade medoid; this score is positive exactly when the genome satisfies the *difficult* criterion previously defined. Per clade, we retain the 5 genomes with the highest scores for the Hard samples, and the 5 with the lowest scores for the Easy samples.

Across all datasets and difficulty levels, we simulate 20 paired-end samples with ART v2016.06.05 [22] (HiSeq 2500 profile, 150 bp reads, 250 bp mean fragment, std 10 bp). Each sample independently draws up to 3 genomes per taxon (without replacement within a sample, and with replacement across samples) from the retained candidates, at uniform per-taxon coverage of approximately 100*×* distributed evenly across drawn genomes.

##### Dirichlet-distributed samples

We evaluate performance under more general randomized compositions by simulating samples with taxon abundances drawn from a symmetric Dirichlet distribution with concentration *α* ∈ *{*0.1, 1.0, 10.0*}*, ranging from sparse compositions with few dominant taxa (*α* = 0.1), to near-equal abundances (*α* = 10.0). For every *α*, we generate 10 compositions each consisting of 20 SARS-CoV-lineages or 5 IAV clades randomly sampled from the available taxa. Per composition, we select up to 10 genomes per taxon by first selecting the medoid, then iteratively adding the genome most distant from all previously selected genomes within the taxon to maximize within-taxon diversity. We then simulate 10 paired-end samples per composition with ART with the parameters described above, at a total per-sample coverage of 2000*×* distributed according to the Dirichlet-sampled taxon abundances. For each taxon, the selected genomes are randomly permuted and assigned to the samples in order, cycling over the permutation when fewer than 10 genomes are available.

#### Profiling pipeline

For all datasets we followed the VLQ pipeline [23] in combination with kallisto v0.44.0 [24] to estimate taxon abundances. For index construction we used default parameters, and we convert output with a custom python script to obtain relative abundances at lineage- and clade-level. Taxa that were reported at an abundance below 0.1% were omitted, and remaining abundances were normalized to sum up to 100%.

#### Evaluation

To evaluate profiling performance across reference sets, we compute per-sample accuracy metrics. For abundance estimation, we use abundance accuracy, ranging from 0 (maximal deviation) to 1 (perfect reconstruction) which is defined as:

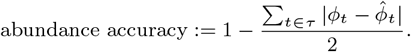

Here, *τ* denotes the set of taxa, *ϕ*_*t*_ the groundtruth abundance of taxon *t*, and 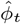 the normalized estimated relative abundance of taxon *t*.

In addition, we compute F1-scores to assess presence/absence detection. A taxon is considered present if its normalized estimated abundance is non-zero. The F1-score is the harmonic mean of precision and recall, ranging from 0 to 1, where 1 indicates perfect identification of present taxa with no false positives, and 0 indicates complete failure (no overlap between predicted and present taxa).

##### Difficulty-stratified evaluation

For each dataset and difficulty, we report per-sample abundance accuracy and F1-score across the 20 samples, summarized as median and interquartile range. To assess whether method differences are statistically significant, we performed paired Wilcoxon signed-rank tests between each ReSeT configuration and the single strongest alternative method, with Benjamini-Hochberg correction applied within per-dataset test families spanning all difficulties, ReSeT configurations and metrics. This strongest alternative is selected per dataset (pooling all difficulty levels and taking medians) by determining the lowest mean rank across both metrics (rank 1 is best).

For both evaluation regimes, when two methods produce identical reference selections, we interpret them as a single combined entry.

#### Parameter tuning

To determine method parameterizations, we perform a parameter tuning step before evaluating profiling performance. Tuning is done independently per virus by creating a temporal train-test split of the reference genomes (see Supplementary Material). We perform a grid search over multiple parameters: for every non-ReSeT selection we consider multiple similarity thresholds and distance estimation methods (MASH v2.3 [14] and sourmash branchwater v0.9.14 [15]) with varying sketch sizes. For ReSeT we search over the selection cost *c* and scale parameter *λ* across all distance estimation settings.

Tuning performance is evaluated on difficulty-stratified samples from held-out reference genomes. Scores are aggregated across difficulty levels using a weighted average that prioritizes Medium and Hard samples over Easy (see Supplementary Material for exact weights), reflecting our interest in performance under increased taxonomic ambiguity. For SARS-CoV-2, we further average weighted scores across the USA and China datasets with equal weight. To select the best parameterization, we consider three tuning targets: F1-tuned (highest weighted mean F1-score), Abundance-tuned (highest weighted mean abundance accuracy) and Rank-tuned (lowest worst-case rank over both metrics), resulting in up to three configurations per method (collapsing configurations resulting in identical reference sets).

## Results

### Taxonomic ambiguity differs across viruses

To characterize the taxonomic ambiguity of our datasets, we quantified the ambiguity of all reference genomes by comparing nearest intra-taxon distances (*d*_intra_) with nearest inter-taxon distances (*d*_inter_) for every genome. In this, we exclude taxa consisting of a single genome, as these will have higher inter-taxon distance by default.

Figure 1**(A)** shows the distribution of log_2_ 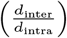 for all three datasets. If this quantity is below 0 for a genome, it means that the genome is closer to a genome of another taxon than any other genome of its own taxon, and we define it as ambiguous. Both SARS-CoV-2 datasets have peaks around 0.3, indicating that overall ambiguity is relatively high. This is further established by the fact that over 9% of the reference genomes are taxonomically ambiguous (i.e. log_2_ 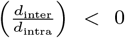). In contrast, the IAV dataset is almost ambiguity free, with the mode of the distribution sitting around 3 and only 0.12% of the genomes being characterized as ambiguous. This two-order-of-magnitude difference in ambiguity between the viruses provides a natural contrast for evaluating whether taxon-aware reference selection improves profiling accuracy.

**Figure 1.**
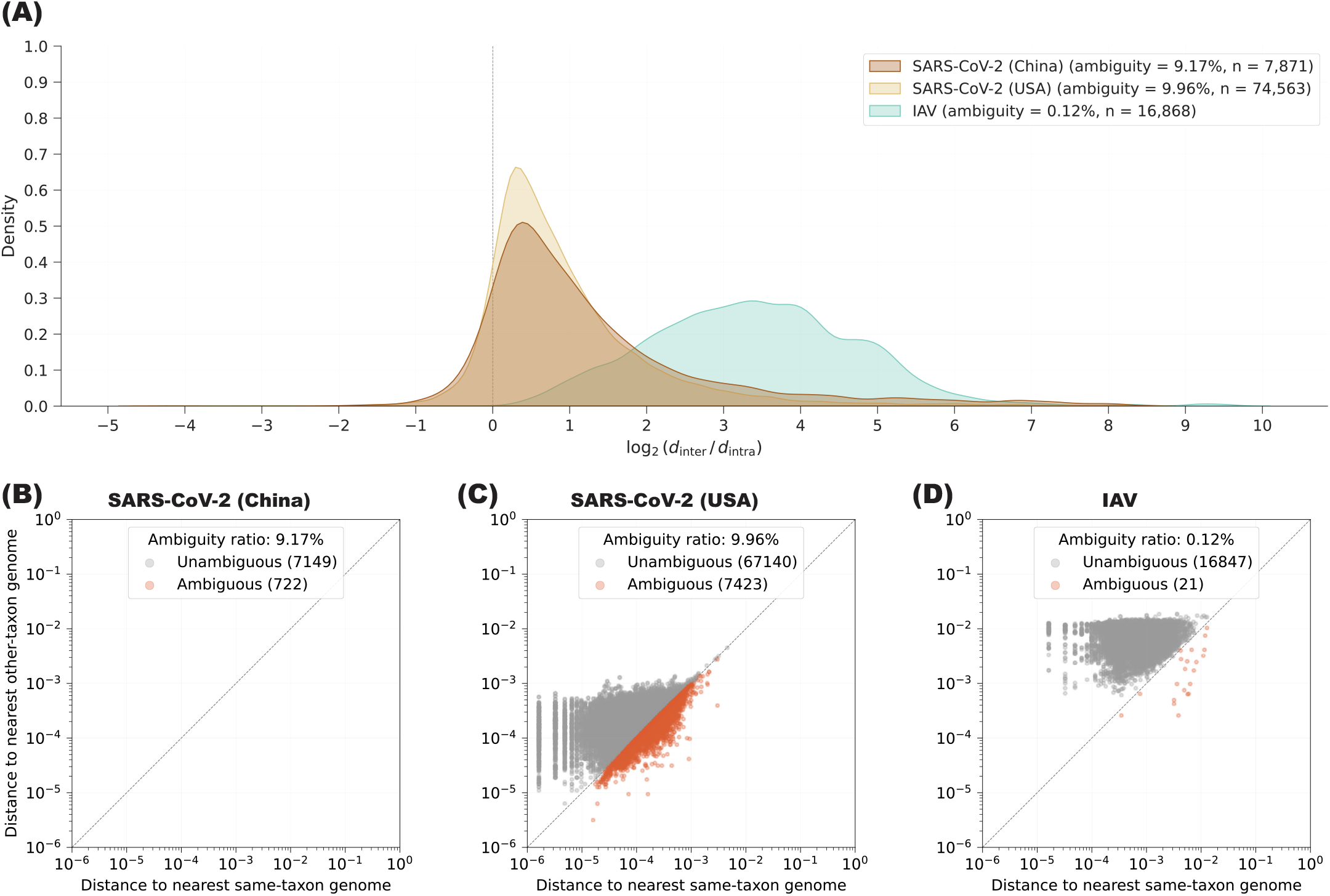

In addition to the distribution, Figures 1**(B)**–**(D)** show the absolute (*d*_intra_, *d*_inter_) pairs in all three datasets. These figures show that distances between genomes are substantially smaller for the SARS-CoV-2 datasets compared to the IAV dataset. Moreover, for the SARS-CoV-2 datasets most of the ambiguous genomes are close to the diagonal that separates ambiguous from unambiguous genomes, whereas ambiguous IAV genomes tend to be further separated. This suggests that SARS-CoV-2 ambiguity is inherently present due to marginal differences between inter- and intra-taxon distances, rather than from isolated outliers as in IAV.

### Parameter tuning results

We selected parameters for ReSeT and all competing methods by a grid search against three tuning targets: F1-score, abundance accuracy and mean combined rank (see Methods). The resulting configurations differ between viruses, and in some cases between tuning targets within the same virus (Supplementary Tables S1–8).

For SARS-CoV-2, both F1- and Abundance-tuned configurations set *λ* = 10^−5^ and *c* = 10^−1^, but differed in distance estimation method: the F1-tuned configuration used MASH with a sketch size of 10,000, and the Abundance-tuned configuration used sourmash (cosine) with a sampling factor of 3. The rank-tuned configuration instead used MASH with a sketch size of 5,000 and set *λ* = 0 and *c* = 1, effectively mimicking a medoid-like selection.

For IAV, one configuration optimized both the F1-score and minimax-rank meaning we only have three distinct configurations. Both configurations used MASH with a sketch size of 500 for distance estimations and set *λ* = 0, but differed in the selection cost parameter *c*, with the F1/rank-tuned configuration setting *c* = 10^−2^ and the Abundance-tuned configuration setting *c* = 1.

Configurations selected for competing methods (medoid, hierarchical clustering, VSEARCH) for both viruses are provided in Supplementary Tables S1–3 (SARS-CoV-2) and Supplmentary Tables S5–7 (IAV).

### ReSeT improves profiling results on stratified difficult samples

To isolate the effect of taxonomic ambiguity, we compared ReSeT to against alternatives on difficulty-stratified samples consisting of either taxonomically ambiguous taxa (Medium and Hard), or taxonomically separable taxa (Easy).

#### SARS-CoV-2 (USA)

ReSeT’s improvements over the strongest alternative (medoid and hierarchical clustering using MASH with a sketch size of 10,000) were strongest on the USA-based SARS-CoV-2 dataset (Figure 2; Supplementary Tables S9–10). On the Hard samples, both the F1-tuned and Rank-tuned configurations significantly improved abundance accuracy (Δ = +0.088 and Δ = +0.093, respectively) and only the F1-tuned configuration significantly improved F1-score (Δ = +0.019), while the Rank-tuned configuration was indistinguishable from the strongest alternative. In contrast, the Abundance-tuned ReSeT configuration showed no significant improvements on abundance accuracy, but significantly degraded F1-score (Δ = −0.041). On the Easy samples, only the F1-tuned configuration significantly outperformed the strongest alternative, improving abundance accuracy (Δ = +0.012) while matching on F1-score. Both the Rank-tuned and Abundance-tuned configurations significantly underperformed on both metrics, with the Abundance-tuned showing the largest losses (Δ_Abund._ = −0.176, Δ_F1_ = −0.083). Results on the Medium samples broadly followed the same trends as Hard samples. From this, it follows that the F1-tuned configuration emerged as the most robust selection on this dataset, combining significant improvements with no significant losses at any difficulty.

**Figure 2.**
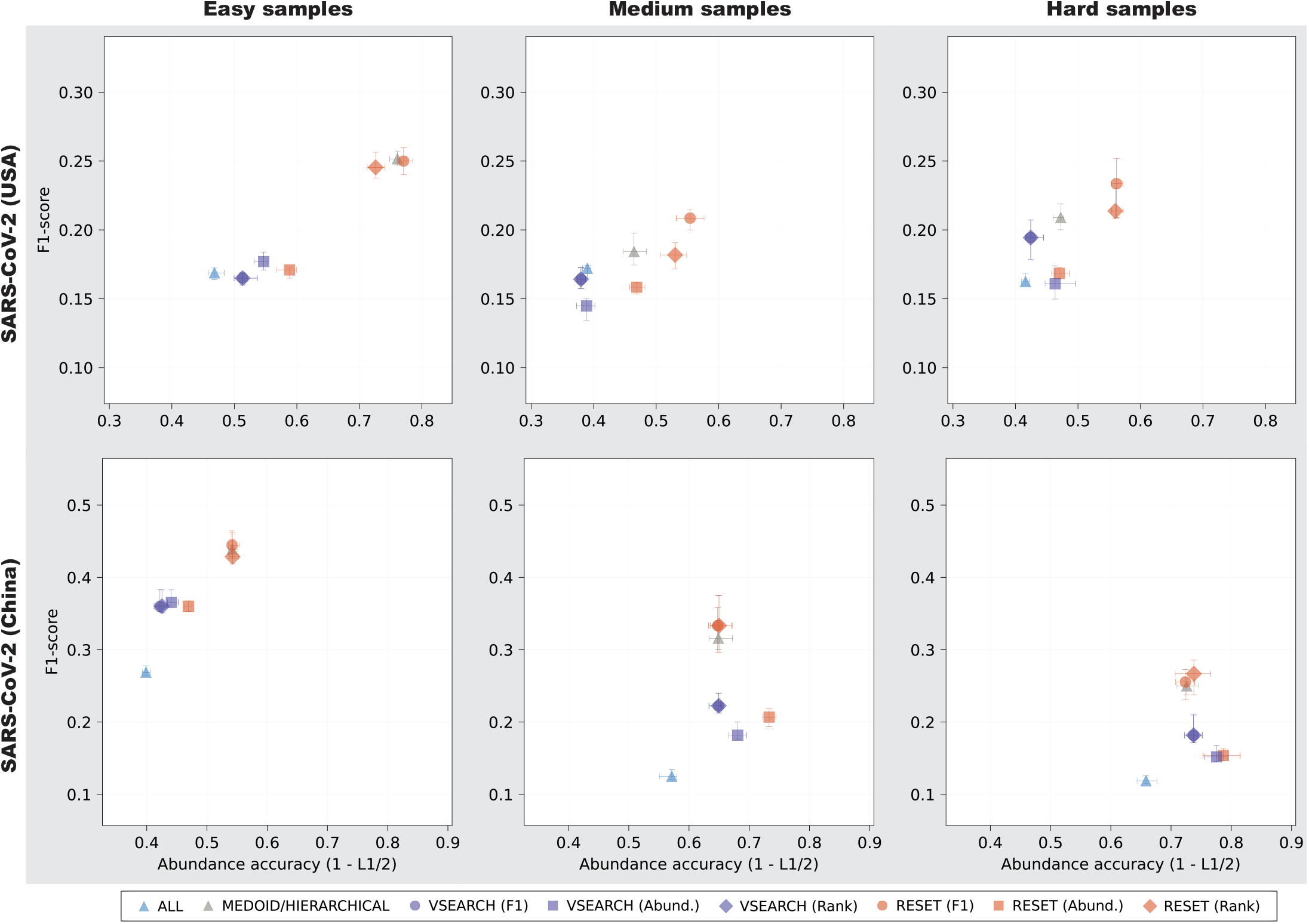
Median abundance accuracy and F1-score across the 20 difficulty-stratified SARS-CoV-2 samples (per difficulty). Error bars indicate IQR ranges, and omission of tuning target means a single configuration was selected for all three targets.

#### SARS-CoV-2 (China)

Improvements over the strongest alternative were less pronounced on the China-based SARS-CoV-2 dataset (Figure 2; Supplementary Tables S9–10), with no single ReSeT configuration universally performing best on both metrics. On the Hard samples, only the Abundance-tuned ReSeT configuration significantly improved abundance accuracy (Δ = +0.066), and only the Rank-tuned configuration significantly improved F1-score (Δ = +0.011). On the Easy samples, no configuration significantly improved over the best alternative, and the Abundance-tuned configuration significantly underperformed on both abundance accuracy (Δ = −0.075) and F1-score (Δ = −0.081). Despite abundance accuracy gains on the Hard samples, this configuration had significantly lower F1-score on the Hard samples (Δ = −0.096). Medium samples again followed the same trends as Hard samples. Overall, the Rank-tuned configuration provided a significant improvement over the strongest alternative without significant losses elsewhere, while the F1-tuned configuration remained statistically indistinguishable from the baseline across all difficulties.

#### IAV

Unlike both SARS-CoV-2 datasets, ReSeT’s performance on the IAV dataset was mixed across difficulties (Figure 3; Supplementary Tables S9–10) with all configurations significantly underperforming the strongest alternative (F1/Rank-tuned VSEARCH at 99% sequence identity) on Easy samples, while significantly improving F1-score on the Hard samples (Δ = +0.190 for both configurations). Despite improvements on F1-score however, abundance accuracy on Hard samples remained significantly below the strongest alternative (Δ_F1/Rank_ = −0.041, Δ_Abund._ = −0.102), indicating that the improved identification of clades comes at the cost of worse clade abundance estimations. Interestingly, F1/Rank-tuned ReSeT’s tends to outperform other alternatives (no-selection, medoid and hierarchical clustering) in terms of F1-score, sometimes at the cost of lower abundance accuracy. Additionally, the strongest VSEARCH configuration was difficulty-dependent, with the F1/Rank-tuned configuration achieving highest abundance accuracy on the Easy samples (both configurations attain perfect F1-score), whereas Abundance-tuned VSEARCH strictly outperformed the F1/Rank-tuned configuration across both metrics on the Hard samples.

**Figure 3.**
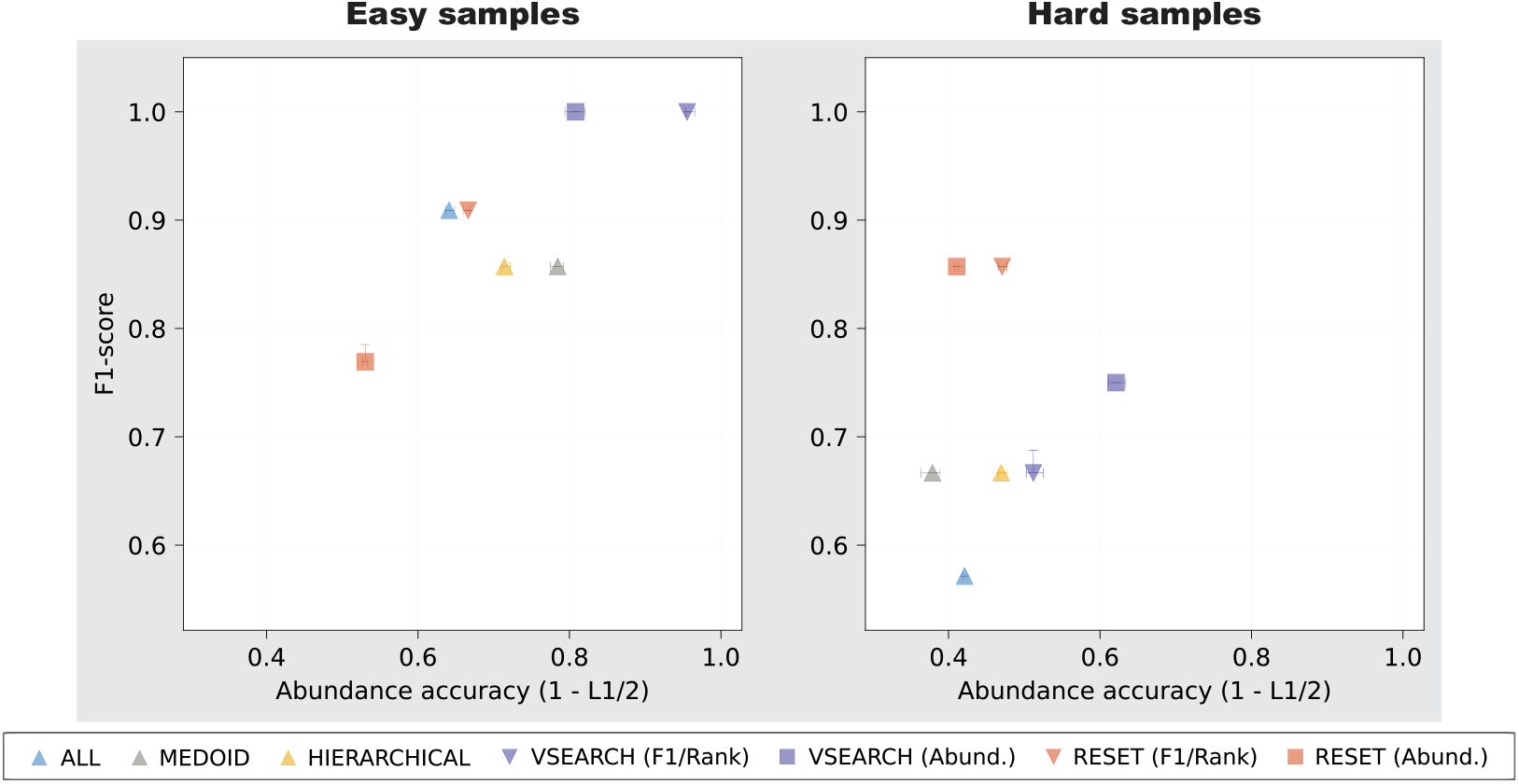
Median abundance accuracy and F1-score across the 20 difficulty-stratified IAV samples (per difficulty). Error bars indicate IQR ranges, and omission of tuning target means a single configuration was selected for all three targets.

### ReSeT performance on random mixtures is regime-dependent

To evaluate ReSeTin a more realistic setting, we compare performance on randomized mixtures sampled from Dirichlet distributions (Table 1). On both SARS-CoV-2 datasets, ReSeT outperformed the alternatives: ReSeT (F1-tuned) achieved the lowest mean rank on the USA-based dataset (avg = 2.78 vs 3.07 for medoid/hierarchical clustering), and ReSeT tuned for Rank on the China-based dataset (avg = 2.13 vs 2.97). ReSeT also achieved sole top rank on a substantially larger share of compositions than the strongest alternative. On the USA-based dataset, F1-tuned ReSeT was the sole top performer on 15/30 F1-score evaluations and 8/30 for abundance accuracy compared to 6/30 and 1/30 respectively for medoid/hierarchical clustering. On the China-based dataset, Rank-tuned ReSeT achieved sole top performance on 18/30 F1-score evaluations and 8/30 for abundance accuracy, compared to 8/30 and 2/30 for medoid/hierarchical clustering.

**Table 1.** Mean ranks and first-place counts for reference selection methods across Dirichlet-distributed compositions (10 compositions *×* 3 concentration parameters = 30 evaluations per dataset). Methods are ranked per composition (rank 1 = best). #1 columns report the number of evaluations where a method achieved the sole top rank; compositions with tied top ranks are not counted. Best value per column in **bold**.

| Method | Mean rank |  |  | #1 (of 30) |  |  |
| --- | --- | --- | --- | --- | --- | --- |
|  | F1 | Abund. | Avg | F1 | Abund. | Avg |
| *SARS-CoV-2 (USA)* |  |  |  |  |  |  |
| All | 7.47 | 7.33 | 7.40 | 0 | 1 | 0 |
| Medoid/Hierarchical clustering | 2.33 | 3.80 | 3.07 | 6 | 1 | 0 |
| VSEARCH (F1) | 4.67 | 4.07 | 4.37 | 0 | 0 | 0 |
| VSEARCH (Abund.) | 5.40 | 4.53 | 4.97 | 0 | 3 | 0 |
| VSEARCH (Rank) | 4.67 | 4.07 | 4.37 | 0 | 0 | 0 |
| ReSeT (F1) | **2.00** | **3.57** | **2.78** | **15** | **8** | **7** |
| ReSeT (Abund.) | 6.63 | 5.03 | 5.83 | 0 | 3 | 0 |
| ReSeT (Rank) | 2.83 | 3.60 | 3.22 | 6 | 5 | 3 |
| *SARS-CoV-2 (China)* |  |  |  |  |  |  |
| All | 7.73 | 4.77 | 6.25 | 0 | 3 | 0 |
| Medoid/Hierarchical clustering | 2.13 | 3.80 | 2.97 | 8 | 2 | 0 |
| VSEARCH (F1) | 6.43 | 6.83 | 6.63 | 0 | 0 | 0 |
| VSEARCH (Abund.) | 4.40 | 3.40 | 3.90 | 1 | **12** | 0 |
| VSEARCH (Rank) | 6.03 | 6.37 | 6.20 | 1 | 0 | 0 |
| ReSeT (F1) | 2.70 | 4.70 | 3.70 | 2 | 0 | 0 |
| ReSeT (Abund.) | 4.93 | 3.50 | 4.22 | 0 | 5 | 0 |
| ReSeT (Rank) | **1.63** | **2.63** | **2.13** | **18** | 8 | **5** |
| *IAV* |  |  |  |  |  |  |
| All | 3.98 | 4.20 | 4.09 | 0 | 3 | 0 |
| Medoid | 4.62 | 3.43 | 4.03 | 1 | 5 | 0 |
| Hierarchical clustering | 4.65 | 3.90 | 4.28 | 0 | 0 | 0 |
| VSEARCH (F1/Rank) | **1.87** | **2.30** | **2.08** | **18** | **14** | **10** |
| VSEARCH (Abund.) | 2.60 | 4.00 | 3.30 | 5 | 1 | 0 |
| ReSeT (F1/Rank) | 3.78 | 3.87 | 3.83 | 2 | 7 | 0 |
| ReSeT (Abund.) | 6.50 | 6.30 | 6.40 | 0 | 0 | 0 |

The optimal ReSeT configuration differed between the two SARS-CoV-2 datasets, with the F1-tuned configuration achieving lowest mean rank on the USA-based dataset, and the Rank-tuned configuration achieving lowest mean rank on the China-based dataset. The Abundance-tuned configuration performed substantially worse on both datasets, with mean ranks of 5.83 (USA) and 4.22 (China) compared to 2.78 and 2.13 for the best performing configuration respectively.

On the IAV dataset, F1/Rank-tuned VSEARCH was consistently dominant, achieving the sole top rank on 18/30 F1-score evaluations and 14/30 for abundance accuracy (10/30 on both metrics simultaneously). The F1/Rank-tuned ReSeT configuration achieved an average rank of 3.83 which is comparable to the medoid and all selections (4.03 and 4.09 resp.), but is well behind VSEARCH (2.08 for F1/Rank-tuned). The Abundance-tuned ReSeT configuration ranked last among all methods on every metric (avg = 6.40), performing worse than even the no-selection baseline.

Results on the Dirichlet-distributed samples were largely consistent with the difficulty-stratified results, with ReSeT outperforming alternatives on SARS-CoV-2 and VSEARCH dominating on IAV. Rankings were stable across all three Dirichlet concentration parameters (Supplementary Table S11), indicating that the relative performance of methods was robust to variation in sample composition. On the SARS-CoV-2 datasets, the no-selection (All) reference set ranked last or near-last, as in the difficulty-stratified experiments. On IAV, however, the All reference set ranked more closely to non-VSEARCH selections, even outperforming the Abundance-tuned ReSeT configuration. Finally, across experiments, the Abundance-tuned ReSeT configuration tended to substantially underperform other ReSeT configurations, suggesting that tuning ReSeT exclusively on abundance accuracy generalizes poorly.

### Parameter tuning reveals cost dominance and scale insensitivity

To characterize the sensitivity of ReSeT to its parameters, we evaluated profiling accuracy across a grid of selection cost *c* and inter-taxon scale *λ* values on the SARS-CoV-2 parameter tuning samples (Supplementary Figures S1–S9). For both SARS-CoV-2 datasets, F1-score improves sharply when *c* increases beyond 10^−3^, with performance plateauing at *c* ≥ 10^−2^. Abundance accuracy shows a similar, but dataset-dependent pattern, with optimal cost being lower for the smaller China dataset (*c* ≈ 10^−4^) than for the larger USA dataset (*c* ≈ 10^−3^).

In contrast, the inter-taxon scale parameter *λ* has minimal effect on both abundance accuracy and F1-score within the recommended cost regime. Across both datasets and metrics, varying lambda from 0 to 10^−1^ produces marginal differences in profiling accuracy, with only *λ* = 1 consistently degrading performance. Based on these findings, it follows that picking *c* ∈ [10^−1^, 10^−3^] and setting *λ* ≈ 10^−5^ works well overall, with *c* potentially requiring adjustments for abundance accuracy-focused applications.

### Computational resource usage

Table 2 shows the resources used by every method across all experiments. ReSeT’s peak memory usage scales quadratically with the total number of input sequences because it relies on a global all-vs-all distance matrix. VSEARCH and both medoid and hierarchical clustering-based approaches are run on a per-taxon basis, which significantly reduces total memory usage for larger datasets (~22.37GB for ReSeT versus ~1.24GB for medoid/hierarchical and ~1.57GB for VSEARCH in the SARS-CoV-2 USA-based dataset). On smaller datasets, such as IAV (16,868 input sequences vs 74,563 for USA), ReSeT’s memory footprint drops to ~2.20GB. The per-taxon methods move in the opposite direction: VSEARCH’s memory increases from *<*1.6GB on USA to ~4.88GB on IAV because peak memory for per-taxon methods scales with the largest taxon, which is larger for IAV (11,716 genomes vs 6,159 in USA).

**Table 2.** End-to-end wall-clock time and peak resident set size (GB) for all selection methods across datasets, using 32 threads. Both time and memory usage include distance computations.

| Method | SARS-CoV-2 (China) |  | SARS-CoV-2 (USA) |  | IAV |  |
| --- | --- | --- | --- | --- | --- | --- |
|  | Time (H:M:S) | RSS (GB) | Time (H:M:S) | RSS (GB) | Time (H:M:S) | RSS (GB) |
| Medoid<sup>1</sup> | 00:05:44 | 0.29 | 00:31:20 | 1.24 | 00:02:23 | 1.05 |
| Hierarchical clustering<sup>1</sup> | 00:05:44 | 0.29 | 00:31:20 | 1.24 | 00:03:33 | 2.61 |
| VSEARCH (F1)<sup>2</sup> | 05:50:17 | 0.17 | 31:57:34 | 0.62 | 00:02:24 | 4.88 |
| VSEARCH (Abund.) | 04:26:25 | 0.47 | 25:03:03 | 1.57 | 00:03:21 | 4.87 |
| VSEARCH (Rank)<sup>2</sup> | 04:16:32 | 0.28 | 25:16:07 | 0.84 | 00:02:24 | 4.88 |
| ReSeT (F1)<sup>3</sup> | 00:24:06 | 1.94 | 19:19:56 | 22.37 | 32:17:50 | 2.20 |
| ReSeT (Abund.) | 07:44:19 | 2.50 | 127:59:07 | 22.36 | 01:14:01 | 2.20 |
| ReSeT (Rank)<sup>3</sup> | 00:10:29 | 1.02 | 05:11:27 | 22.37 | 32:17:50 | 2.20 |

<sup>1</sup>For SARS-CoV-2, medoid and hierarchical selections are identical.
<sup>2</sup>For IAV, F1- and Rank-tuned VSEARCH yielded identical parameterizations.
<sup>3</sup>For IAV, F1- and Rank-tuned ReSeT yielded identical parameterizations.

ReSeT’s runtime appears governed by three factors that collectively determine neighborhood sizes: the selection cost parameter (higher costs select fewer genomes and finish faster), the choice of distance estimation method (MASH is substantially cheaper than sourmash), and the number of genomes per taxon. The Abundance-tuned SARS-CoV-2 (USA) run, which uses sourmash-based cosine similarity, is the worst case at 128 hours of wall-clock time. MASH-based configurations on the same dataset complete in 5–19 hours, and on the smaller China-based dataset they finish in under 25 minutes. IAV illustrates the taxon-density extreme: its largest clade drives F1/Rank-tuned ReSeT runs to more than 32 hours, while the Abundance-tuned configuration completes in approximately an hour because it omits the inter-taxon term and has a high selection cost. The baseline methods all operate on a per-taxon basis, parallelize trivially across taxa, and finish selection for the largest single-taxon within an hour. ReSeT is therefore practical when taxa contain on the order of thousands of genomes and MASH-based distances suffice. For taxa exceeding ~10,000 genomes, only high-cost configurations remain tractable.

## Discussion

We designed ReSeT as a facility-location-based reference genome selection tool that operates on arbitrary distance matrices and incorporates tunable inter-taxon discrimination to improve taxonomic profiling accuracy. Across both evaluation frameworks (difficulty-stratified and Dirichlet-distributed compositions), ReSeT provided its most consistent improvements on the high-ambiguity SARS-CoV-2 datasets, where it outperformed or matched the strongest baselines depending on configuration. On the low-ambiguity IAV dataset, VSEARCH tuned for Rank/F1-score remained dominant, although ReSeT improved F1-score on the Hard samples. These results indicate that the benefits of different selection mechanisms are strongly regime-dependent. ReSeT improves accuracy on datasets that exhibit high taxonomic ambiguity, often competing with medoid/hierarchical clustering-based selection, while underperforming VSEARCH, but performing comparably against other methods in datasets where this ambiguity is missing.

ReSeT occupies a practical intermediate between existing facility-location-based approaches for reference genome selection. PARNAS [11] solves a related problem, but requires a tree metric, limiting applicability where reliable phylogenies are unavailable. Repset [12] achieves an optimal submodular approximation guarantee, but is designed for protein sequences and currently relies on PSI-BLAST [13] for distance computations, which is prohibitive at the genome scales common in taxonomic profiling. ReSeT instead accepts general pairwise distance matrices, relaxing these constraints at the cost of losing optimality guarantees.

The performance gap between ReSeT and traditional approaches appears strongly dependent on the taxonomic ambiguity of the reference database. We quantified this through the ambiguity index (i.e. the fraction of genomes for which *d*_inter_ *< d*_intra_), which is computable from pairwise distances and taxon labels alone. In our datasets, this index characterizes two distinct regimes: high-ambiguity (SARS-CoV-2), where *>*9% of the reference genomes are classified as ambiguous, and where, on average, genomes are only 2*×* closer to their own lineage than others, versus low-ambiguity (IAV) where less than 0.13% of genomes are considered ambiguous and genomes are 10*×* closer to their own clade on average. ReSeT’s advantage follows this split directly (dominating on SARS-CoV-2, absent on IAV) and remains consistent across both evaluation regimes and all Dirichlet concentrations, suggesting that ambiguity, rather than sample composition, is the primary driver of relative performance. As the index depends only on information required to run ReSeT, it functions as a practical pre-screening criterion: users can estimate whether facility-location-based selection is likely to outperform more traditional methods before committing to reference set construction. However, our experiments span only two viruses with ambiguity indices differing by two orders of magnitude, leaving the threshold at which facility-location-based selection becomes advantageous unresolved. Aggregating SARS-CoV-2 lineages at coarser taxonomic levels would generate datasets with intermediate ambiguity, allowing this threshold to be characterized directly.

To our knowledge, ReSeT is the first reference genome selection method to incorporate inter-taxon discrimination as an explicit, tunable term in the selection objective. While ReSeT produces favorable reference sets, our parameter tuning results show that the inter-taxon distance term contributes only marginally, even in the high-ambiguity regime where discriminative selection is expected to matter most. The discriminative information gained from *f*_inter_ may be redundant, with intra-taxon coverage and cost control being sufficient for optimizing profiling accuracy. Alternatively, our max-similarity formulation may be too limited to meaningfully incorporate inter-taxon discrimination, as it reduces each taxon pair to its single closest contact and discards broader overlap patterns. Systematic comparisons of alternative inter-taxon penalty definitions (e.g. mean, top-*k*, or soft-max similarity across taxon pairs) could help distinguish between these cases by testing whether more permissive formulations capture meaningful patterns of taxon entanglement that max-similarity cannot. Such formulations would sacrifice our linear problem structure, but may reveal whether *f*_inter_’s limited contribution is due to redundancy or inadequacy.

ReSeT relies on an all-vs-all pairwise distance matrix, which limits practical scalability to databases of ~ 10^5^ genomes. One natural consequence is that ReSeT’s resource usage is substantially larger than per-taxon baselines: since memory usage scales quadratically with the number of candidate reference genomes, the peak memory usage reaches approximately 23GB on the USA-based SARS-CoV-2 dataset (74,563 genomes), compared to *<*2GB for medoid, hierarchical clustering and VSEARCH selections. Runtime follows a similar pattern; the F1-optimized ReSeT selection requires approximately 15 hours of wall-clock time on the same dataset, whereas VSEARCH achieves comparable aggregate runtimes, but is trivially parallelizable across taxa. These computational constraints are an inherent consequence of global, rather than per-taxon optimization. Per-taxon baselines avoid this cost by partitioning the problem along taxonomic boundaries, which keeps their distance matrices local but omits inter-taxon information entirely. An intermediate strategy would partition the database via efficient coarse clustering tools such as LINCLUST [25] (similar to dRep’s approach [9]), reducing the all-vs-all footprint while preserving the ability to incorporate confusable taxa jointly.

An important future direction is to investigate the generalizability of the results in this work. First, all evaluations relied on a single profiling pipeline (VLQ [23]), and assessing whether outcomes are consistent across alternative pipelines, particularly those using different read-assignment strategies (e.g. [26, 27]), would establish whether ReSeT’s gains are pipeline-specific or apply more broadly. Second, the optimal ReSeT configuration differs across datasets we evaluated, suggesting that parameter tuning may not transfer directly without re-optimization. Further characterizing how dataset properties such as scale and composition shape the parameter landscape would offer guidance on how to configure ReSeT for new datasets. Finally, our experiments are limited to viral datasets with short genomes and high mutation rates. Extending the scope to bacterial communities, for example through established benchmarks such as CAMI [28, 29], would test whether ReSeT’s merits generalize beyond viral datasets, in settings where intra- and inter-taxon diversity can be structured differently.

Overall, our results show that ReSeT-based reference genome selection is most useful in regimes where taxa are not cleanly separated, while offering limited advantage when they are. Our proposed ambiguity index further provides a concrete diagnostic for this distinction: users can compute it from pairwise distances already required to run ReSeTto determine whether its increased computational cost is justified. This makes ReSeT particularly suited to settings where high-resolution profiling is required under weak taxonomic separation, such as SARS-CoV-2 lineage abundance estimation in wastewater-based epidemiology [23, 4], strain-resolution profiling in microbiome studies [2], or potentially sub-species profiling of bacterial pathogens where clinically relevant distinctions occur below the species level.

## Conflicts of interest

The authors declare that they have no competing interests.

## Funding

J.A.B. is supported by the research programme Veni (file number VI.Veni.242.338), which is financed by the Dutch Research Council (NWO).

## Data availability

The datasets used for the experiments in this work are available on Zenodo at https://doi.org/10.5281/zenodo.20553987

## Author contributions

J.B. and J.A.B. conceived the study. J.B. developed the method with feedback from J.A.B. J.B. implemented the method and performed data analysis with feedback from J.A.B. J.B. and J.A.B. wrote the manuscript. All authors read and approved the final manuscript.

## Acknowledgements

We gratefully acknowledge all data contributors, i.e., the Authors and their Originating laboratories responsible for obtaining the specimens, and their submitting laboratories for generating the genetic sequence and metadata and sharing via the GISAID Initiative, on which part of this research is based.

Research reported in this work was partially or completely facilitated by computational resources and support of the Delft AI Cluster (DAIC) at TU Delft (RRID: SCR 025091), but remains the sole responsibility of the authors, not the DAIC team.

## Supplementary Information

### Supplementary Material

Includes Supplementary Information, all Supplementary Figures and Supplementary Tables.

## Notes

https://doi.org/10.5281/zenodo.20553987

https://github.com/JaspervB-tud/ReSeT

### Competing Interest Statement

The authors have declared no competing interest.

