## Supplementary Material for "ReSeT: a taxonomy-aware reference genome selection tool"

June 17, 2026

### 1 ReSeT

As stated in the Methods section, we solve a Facility Location Problem (FLP) based model which simultaneously optimizes the representation of and discriminability between taxa. Here we will again provide all of the necessary details for the model, after which we provide the complete problem formulated as a mixed integer linear program (MILP).

#### 1.1 Notation

Let $\mathcal{G}$ denote a set of candidate reference genome assemblies, $T$ a set of taxa and $\tau : \mathcal{G} \to T$ a function that maps every genome in $\mathcal{G}$ to a single taxon in $T$. We further assume a distance function $d : \mathcal{G} \times \mathcal{G} \to [0, 1]$ which is symmetric ($d(g, g') = d(g', g)$) and satisfies $d(g, g) = 0$, without having to satisfy additional properties. This set-up is flexible enough to be compatible with most practical sequence distance/similarity estimators, which generally output genetic distances as a (dis)similarity which naturally satisfies the range condition. For convenience, we let $\mathcal{G}_t := \{g \in \mathcal{G} : \tau(g) = t\}$ denote the set of genomes from taxon $t \in T$ and for any $G \subseteq \mathcal{G}$ we similarly define $G_t := \{g \in G : \tau(g) = t\}$. Finally, we assume an arbitrary total order on both $T$ and $\mathcal{G}$ so that $\{t \in T : t < t'\}$ denotes all taxa preceding $t'$ and $\{g \in \mathcal{G} : g < g'\}$ denotes all genomes that precede $g'$.

#### 1.2 Decision variables.

To model the problem as a MILP, we now introduce the following primary decision variables:

$$x_g = \begin{cases} 1, & \text{if genome } g \text{ is selected,} \\ 0, & \text{otherwise.} \end{cases} \qquad \forall g \in \mathcal{G}$$

$$y_{g,g'} = \begin{cases} 1, & \text{if genome } g' \text{ is selected as a representative for } g, \\ 0, & \text{otherwise.} \end{cases} \qquad \forall g, g' \in \mathcal{G} : \tau(g) = \tau(g')$$

These variables are central to our FLP formulation, and model which genomes are selected, and which genomes are represented by which genomes. In addition to the above decision variable, we also introduce the following auxiliary variables:

$$z_{g,g'} = \begin{cases} 1, & \text{if genome } g \text{ and } g' \text{ are both selected,} \\ 0, & \text{otherwise.} \end{cases} \qquad \forall g < g' \in \mathcal{G} : \tau(g) < \tau(g')$$

These variables will be used to model ReSeT's novel inter-taxon penalization, but break the submodularity of the selection objective as it no longer allows us to solve the problem on a per-taxon basis.

#### 1.3 Constraints.

Our model includes the following constraints:

$$\sum_{g' \in \mathcal{G}_{\tau(g)}} y_{g,g'} = 1 \qquad \forall g \in \mathcal{G}, \tag{1}$$

$$y_{g,g'} \leq x_{g'} \qquad \forall g, g' \in \mathcal{G} : \tau(g) = \tau(g'), \tag{2}$$

$$z_{g,g'} \geq x_g + x_{g'} - 1 \qquad \forall g < g' \in \mathcal{G} : \tau(g) < \tau(g'), \tag{3}$$

$$z_{g,g'} \leq 0.5(x_g + x_{g'}) \qquad \forall g < g' \in \mathcal{G} : \tau(g) < \tau(g'). \tag{4}$$

Here, constraints (1)–(2) together force that every genome $g \in \mathcal{G}$ needs to be represented by either itself ($x_g = 1$) or some other genome of the same taxon ($x_g = 0$ and $y_{g,g'} = 1$ for some $g' \in \mathcal{G} \setminus \{g\}$ : $\tau(g) = \tau(g')$), implicitly ensuring that every taxon is represented. Furthermore, constraints (3)–(4) force $z_{g,g'}$ to one if both $x_g$ and $x_{g'}$ are set to 1 (that is, both $g$ and $g'$ are selected). Note that in practice, since we are minimizing our objective, (4) are redundant and can be omitted.

In addition to the variables and constraints introduced so far, we also introduce the auxiliary variables $q_{t,t'}$ $\forall t, t' \in T : t < t'$ which represent the shortest distance (highest similarity) across taxon pairs. To enforce that $q_{t,t'}$ takes this value, we include the following constraints:

$$q_{t,t'} \geq (1 - d(g,g'))\, z_{g,g'} \qquad \forall t < t',\ \forall g < g' : \tau(g) = t\ \tau(g') = t' \tag{5}$$

which, through (3) and the fact that we are minimizing our objective, set $q_{t,t'}$ to the highest similarity between any pair of selected genomes from $t$ and $t'$.

##### 1.4 Objective function components.

The objective function of our model now consists of the three components (selection cost, intra-taxon coverage and inter-taxon separation) described in the Methods section of the main paper. Reformulating to conform with our definitions so far this yields:

$$\min\, c \sum_{g \in \mathcal{G}} x_g + \sum_{\substack{g,g' \in \mathcal{G} \\ \tau(g)=\tau(g')}} d(g,g') y_{g,g'} + \lambda \frac{2(|\mathcal{G}| - |T|)}{|T|(|T|-1)} \sum_{t<t'} q_{t,t'}.$$

Here we implicitly assume that $|T| \geq 2$, as otherwise the normalization is ill-defined. However, when $|T| = 1$, the inter-taxon penalization disappears entirely as there exist no distinct $t < t'$, making the model well-defined either way.

##### 1.5 Full model specification.

Combining all of the components introduced here (except the redundant constraints in (4)), we obtain the full model specification below:

$$\begin{aligned}
\min \quad & c \sum_{g \in \mathcal{G}} x_g + \sum_{\substack{g,g' \in \mathcal{G} \\ \tau(g)=\tau(g')}} d(g,g') y_{g,g'} + \lambda \frac{2(|\mathcal{G}| - |T|)}{|T|(|T|-1)} \sum_{t<t'} q_{t,t'} \\
\text{s.t.} \quad & \sum_{g' \in \mathcal{G}_{\tau(g)}} y_{g,g'} = 1 & \forall g \in \mathcal{G},
\end{aligned}$$

$$\begin{aligned}
y_{g,g'} &\leq x_{g'} & & \forall g, g' \in \mathcal{G} : \tau(g) = \tau(g'), \\
z_{g,g'} &\geq x_g + x_{g'} - 1 & & \forall g, g' \in \mathcal{G} : g < g',\ \tau(g) < \tau(g'), \\
q_{t,t'} &\geq (1 - d(g,g')) z_{g,g'} & & \forall t, t' \in T : t < t',\ \forall g, g' \in \mathcal{G} : g < g',\ \tau(g) = t,\ \tau(g') = t', \\
x_g &\in \{0,1\} & & \forall g \in \mathcal{G}, \\
y_{g,g'} &\in \{0,1\} & & \forall g, g' \in \mathcal{G} : \tau(g) = \tau(g'), \\
z_{g,g'} &\in \{0,1\} & & \forall g, g' \in \mathcal{G} : g < g',\ \tau(g) < \tau(g'), \\
q_{t,t'} &\geq 0 & & \forall t, t' \in T : t < t'.
\end{aligned}$$

#### 2 Theoretical hardness of ReSeT's selection problem

We will prove the theoretical complexity of ReSeT's selection problem by proving hardness through a reduction from the Set Cover Problem (SCP) with unit set costs, which has been shown to be NP-complete [2]. As completeness is usually restricted to decision problems rather than optimization problems due to requiring membership in NP, we will first formally define two related formulations of our problem:

**Problem 1** (ReSeT-Dec)**.** Given a set of candidate reference genomes $\mathcal{G}$, a set of taxonomic labels $T$ along with a taxonomic labeling function $\tau : \mathcal{G} \to T$, a distance function $d : \mathcal{G} \times \mathcal{G} \to [0,1]$, a selection cost $c$, a inter-taxon scaling factor $\lambda \in \mathbb{R}_{\geq 0}$ and a threshold $K \in \mathbb{Q}_{\geq 0}$, does there exist a feasible selection $G \subseteq \mathcal{G}$ with objective value smaller or equal to $K$?

**Problem 2** (ReSeT-Opt))**.** Given a set of candidate reference genomes $\mathcal{G}$, a set of taxonomic labels $T$ along with a taxonomic labeling function $\tau : \mathcal{G} \to T$, a distance function $d : \mathcal{G} \times \mathcal{G} \to [0,1]$, a selection cost $c$, a inter-taxon scaling factor $\lambda \in \mathbb{R}_{\geq 0}$, find a feasible selection $G \subseteq \mathcal{G}$ that minimizes the objective value.

From this definition, it is clear that ReSeT-Dec is in NP, as a selection $G \subseteq \mathcal{G}$ can be checked to be feasible by scanning all $g \in G$, and given $G$, the variables $(x, y, z, q)$ can be computed in polynomial time, so that the objective value is computable in polynomial time.

As we will reduce from the decision version of the SCP, we provide it below for completeness:

**Problem 3** (SCP-Dec)**.** Given a universe of elements $\mathcal{U} = \{e_1, \ldots, e_n\}$, a collection $\mathcal{S} = \{S_1, \ldots, S_m\}$ of subsets of $\mathcal{U}$ such that $\bigcup_{S \in \mathcal{S}} S = \mathcal{U}$, and an integer $K \in \mathbb{N}$, does there exist $\mathcal{C} \subseteq \mathcal{S}$ such that $|\mathcal{C}| \leq K$ and $\bigcup_{S \in \mathcal{C}} S = \mathcal{U}$?

#### 2.1 Instance construction

Here we describe how to obtain an instance of ReSeT-Dec from an instance of SCP-Dec. First, we fix an integer $l \geq 0$, which will represent the number of taxa $|T|$ minus 1. Then, given an instance of SCP-Dec $(\mathcal{U}, \mathcal{S}, K)$, we construct a ReSeT-Dec as follows.

**Genomes and taxa.** Let

$$\mathcal{G} = \{g_1, \ldots, g_m\} \cup \{h_1, \ldots, h_n\} \cup \{a_1, \ldots, a_l\},$$

where every genome $g_j$ represents a set $S_j \in \mathcal{S}$ and every genome $h_i$ represents an element $e_i \in \mathcal{U}$. Additionally, the $a_k$ are auxiliary genomes added to model the existence of multiple taxa by setting $\tau(g_j) = \tau(h_i) = 0$ for all $j, i$, and $\tau(a_k) = t_k$ for $k = 1, \ldots, l$ such that $|T| = l + 1$. Note that setting $l = 0$ still maintains a feasible construction with no auxiliary taxa, so that the proof below will generalize to any number of taxa.

**Distances.** To model the distances used in ReSeT-Dec, we define $d : \mathcal{G} \times \mathcal{G} \to [0, 1]$ as follows:

$$d(g, g') = \begin{cases} 0 & \text{if } g, g' \in \{g_1, \ldots, g_m\}, \\ 0 & \text{if } \{g, g'\} = \{g_j, h_i\} \text{ and } e_i \in S_j, \\ 0 & \text{if } g = g', \\ 1 & \text{otherwise.} \end{cases}$$

This sets the distance between "set-genomes" equal to 0, and it further sets the distance between an "element-genome" and any "set-genome" that contains it to 0, allowing for free representation in these cases. All other cases where $g \neq g'$ set the distance to 1, including distances to the auxiliary genomes $a_k$.

**Parameters.** The remaining parameters consist of the per-genome selection cost $c$, which we set to 1, and the inter-taxon scaling factor $\lambda$ which we allow to take any arbitrary value. Finally, if $K$ was the threshold parameter used in SCP-Dec, we take $K + l$ as threshold in ReSeT-Dec to account for the additional taxa $t_1, \ldots, t_l$.

This construction is easily seen to be polynomial (both in space and time) in $|\mathcal{U}|$ and $|\mathcal{S}|$.

#### 2.2 Construction properties

With the construction described above, we can obtain an instance of ReSeT-Dec from any SCP-Dec instance. This construction also satisfies the following three properties:

(P1): *Any feasible solution $G$ to the constructed ReSeT-Dec instance contains all $a_k$.* This is a direct consequence of every $a_k$ being the sole representatives of their taxa, further leading to a guaranteed selection cost of at least $c \cdot l = l$.

(P2): *The inter-taxon penalty vanishes.* For any $t < t'$ and any pair $(g, g')$ with $\tau(g) = t$, $\tau(g') = t'$, we have $d(g, g') = 1$ by construction. Hence $1 - d(g, g') = 0$ for any feasible solution, and the value of $\lambda$ does not affect the objective.

(P3): *At least one genome in $\{g_1, \ldots, g_m, h_1, \ldots, h_n\}$ is selected by any feasible solution $G$.* This follows directly from the fact that these are exactly the representatives of taxon $t_0$, and thus by constraints (1) at least one of these must be selected.

#### 2.3 "Swapping lemma"

Our NP-completeness proof will rely on the following lemma:

**Lemma.** *Given any feasible solution $G \subseteq \mathcal{G}$ with objective value $f(G)$ and $G \cap \{h_1, \dots, h_n\} \neq \emptyset$, we can obtain another feasible solution $G'$ such that $G' \cap \{h_1, \dots, h_n\} = \emptyset$ and $f(G') \leq f(G)$.*

The main consequence of this lemma is that we can always transform a feasible solution that selects element-genomes into an equivalent or better feasible solution with no such genomes selected. The proof is given below:

*Proof.* Let $G$ be a feasible solution with $G \cap \{h_1, \dots, h_n\} \neq \emptyset$. We can assume without loss of generality that $h_1 \in G$, which together with the condition that $\bigcup_{S \in \mathcal{S}} = \mathcal{U}$ implies that there exists at least one set $S_j$ such that $e_1 \in S_j$. By construction, we have that $d(h_i, h_{i'}) = 1$ for all $i \neq i'$, such that the minimal distance of $h_1$ to any other genome is 1, except to set-genomes $g_{j'}$ for which $e_1 \in S_{j'}$. Based on the existence of such an $S_j$, we can thus differentiate between two cases: either there exist some $S_j \in \mathcal{S}$ such that $e_1 \in S_j$ and $g_j \in G$, or no such $S_j$ exists.

In the first case, we know through (P1) that $\{h_1, g_j, a_1, \dots, a_l\} \subseteq G$, which means that removing $h_1$ from $G$ to obtain $G'$ maintains feasibility as every taxon remains represented. Furthermore, as $d(h_1, g_j) = 0$, it follows that removing $h_1$ from $G$ reduces the total selection cost by 1 while not the cost of having to represent $h_1$. Finally, as $d(g_j, g_{j'}) = 0$ for every $j'$ and $d(h_1, h_i) = 1$ for every $i \geq 2$, it follows that removing $e_1$ from $G$ does not affect $f_{\text{intra}}$. Thus, we conclude that removing $h_1$ from $G$ maintains feasibility, and doing so reduces the objective function value by exactly 1 meaning that $f(G') < f(G)$.

In the second case, we have that there exists no $g_j \in G$ such that $e_1 \in S_j$. Assuming, without loss of generality, that $e_1 \in S_1$, this means that we can add $g_1$ to $G$ while removing $h_1$ to obtain $G'$. Doing so does not affect feasibility since every taxon remains represented, and as $d(h_1, h_i) = d(h_1, g_j) = 1 \geq d(h_i, g_1)$ for every $i \geq 2$ and $g_j \in G$ it follows that $f(G') \leq f(G)$ (with the inequality being strict if another element-genome previously covered by $h_1$ is now covered by $g_1$ at cost 0), meaning that this never increases the objective function value.

As we can repeatedly apply the removal and swapping processes described above to replace all element-genomes, it inductively follows that doing so yields a feasible solution $G''$ with $G'' \cap \{h_1, \dots, h_n\} = \emptyset$ and $f(G'') \leq f(G)$. □

This lemma immediately provides the following corollary:

**Corollary.** *Any feasible optimal solution $G^*$ can be transformed in an equivalent feasible optimal solution that contains no element-genomes.*

#### 2.4 Main result: hardness proof

**Theorem.** *ReSeT-Dec is NP-complete.*

*Proof.* We have already established membership in NP above. For NP-completeness, we will show that given $K \in \mathbb{N}$, the constructed ReSeT-Dec admits a feasible solution with objective value at most $K + l$ if and only if the original SCP-Dec instance admits a set cover of size at most $K$.

($\Rightarrow$) Assume the constructed ReSeT-Dec instance has a feasible solution $G$ with objective at most $K + l$. Then, by virtue of our swapping lemma and (P3), we know that there exists an equivalent feasible solution $G'$ (possibly the same) which contains none of the element-genomes. Thus, without loss of generality, we can assume that $G$ only consists of elements from $\{g_1, \dots, g_m\}$ and the full collection $\{a_1, \dots, a_k\}$ (P1). From this, (P2), and the fact that $c = 1$, it follows that:

$$f(G) = l + \underbrace{|G \setminus \{a_1, \dots, a_l\}|}_{=|\{g_j \in G\}|} + \underbrace{\sum_{g \in \mathcal{G} \setminus G} \min_{g_j \in G} d(g_j, g')}_{=\sum_{h_i} \min_{g_j \in G} d(g_j, h_i)}$$

This shows that the objective value is equal to: the number of auxiliary taxa $l$, the number of selected set-genomes $|\{g_j \in G\}|$ and the number of element-genomes which are represented by a selected set-genome, for which the corresponding set does not contain the corresponding element. As $f(G) \leq K + l$, this furthermore shows that $|\{g_j \in G\}| + |\{h_i : \nexists S_j \text{ such that } g_j \in G \text{ and } e_i \in S_j\}| \leq K$. Reconsidering the original SCP-Dec formulation, this means that the cost is equal to a number of sets plus a number of elements not covered by the selected sets. As every element occurs in at least one set, we could, for every uncovered element, simply add an arbitrary unselected set that contains it, which shows that the total number of selected sets in such a set cover would contain at most $K$ elements.

($\Leftarrow$) Assume now that SCP-DEC admits a feasible solution with objective value at most $K$. This means that there is a selection of sets $C \subseteq \mathcal{S}$ such that $|C| = K$ and $\bigcup_{S \in C} S = \mathcal{U}$. By construction, selecting exactly those set-genomes in the corresponding RESET-DEC instance means that the total intra-taxon distance is zero, since every element-genome and every set-genome is "covered" by a selected set-genome. To now make this solution feasible, we only need to include the $l$ auxiliary genomes, which yields a feasible solution $G$ consisting of at most $K$ set-genomes (at a cost of $c = 1$ per genome) which represent all element-genomes, and the $l$ auxiliary genomes. Thus we have constructed a feasible RESET-DEC solution with cost at most $K + l$. □

The NP-completeness of RESET-DEC immediately implies NP-hardness of RESET-OPT:

**Corollary.** *RESET-OPT is NP-hard.*

### 3 Size of neighborhoods during local-search

Given an instance of RESET-OPT $(\mathcal{G}, T, \tau, c, \lambda)$ and a feasible solution $G \subseteq \mathcal{G}$, we can derive the exact size of the neigborhood of $G$ under the moves defined in the main paper (add, remove, swaps). Throughout we will use $\overline{G} = \mathcal{G} \setminus G$ to denote the complement of a solution $G$ with respect to $\mathcal{G}$, and we let $N := |\mathcal{G}|$ and $N_t := |\mathcal{G}_t|$.

**Add-moves.** Add-moves will never make a solution infeasible, and can independently be performed for every taxon $t \in T$. Therefore, provided a solution $G \subseteq \mathcal{G}$, there are $|\overline{G}_t|$ possible add-moves from $G$. Combining this for all taxa yields:

$$\sum_{t \in T} |\overline{G}_t| = \sum_{t \in T} (|\mathcal{G}_t| - |G_t|) \leq \sum_{t \in T} |\mathcal{G}_t| = |\mathcal{G}|. \tag{6}$$

Thus the exact number of add-moves is $\sum_{t \in T} |\overline{G}_t|$ which is trivially upperbounded by the total number of candidate genomes $|\mathcal{G}|$ which is of order $O(N)$.

**Remove-moves.** Unlike add-moves, remove-moves can violate feasibility is a taxon is represented by a single candidate reference. Thus we can do something analogue to before, only restricting to taxa with at least two representatives:

$$\sum_{\substack{t \in T \\ |G_t| \geq 2}} |G_t| \leq \sum_{t \in T} |G_t| \leq \sum_{t \in T} |\mathcal{G}_t| = |\mathcal{G}|. \tag{7}$$

It should be understood, however, that add- and remove-moves are complementary (which can be seen from the fact that (6) sums $|\overline{G}_t|$, and (7) sums $|G_t|$). Consequently, as the number of remove-moves is of order $O(N)$ too, we have that the total number of add- and remove-moves is collectively of order $O(N)$.

**Swap-moves.** We will begin our analysis by considering the more general scenario where $m$-swaps are allowed for arbitrary $m$, and we will conclude with $m = 2$ as a special case. Unlike remove-moves, swap-moves will always maintain feasibility as they are performed within taxa and not across. First we consider the number of ways in which we can add $m$ unselected genomes from a taxon $t \in T$:

$$\sum_{k=1}^{\min\{m, |\overline{G}_t|\}} \binom{|\overline{G}_t|}{k}. \tag{8}$$

For simplicity we will denote $m_t := \min\{m, |\overline{G}_t|\}$. Since every swap-move further requires the removal of exactly one selected genome, this means that the total number of swap-moves (allowing up to $m$ additions) within $t$ is equal to:

$$|G_t| \sum_{k=1}^{m_t} \binom{|\overline{G}_t|}{k}. \tag{9}$$

It can easily be seen now that if $m \geq |\overline{G}_t|$ (i.e. $m_t = |\overline{G}_t|$) we have:

$$|G_t| \sum_{k=1}^{m_t} \binom{|\overline{G}_t|}{k} = |G_t| \sum_{k=1}^{|\overline{G}_t|} \binom{|\overline{G}_t|}{k} = |G_t| \left(2^{|\overline{G}_t|} - 1\right). \tag{10}$$

Thus, if $m < |\overline{G}_t|$, we have that the total number of swap-moves within a taxon $t$ is of order $O(N \cdot N^m)$, as $|G_t| \leq |\mathcal{G}_t|$ and $m_t \leq |\overline{\mathcal{G}}_t| \leq |\mathcal{G}_t|$. If $m \geq |\overline{G}_t|$, this instead turns into $O(N \cdot 2^N)$ by (10). This result is exactly what motivates our choice of setting $m = 2$, as then the number of swap-moves for a given taxon is of order $O(N^3)$, yielding a total of $O(|T| \cdot N^3)$ over all taxa.

**Total.** Combining everything, we find that the total number of permitted moves for a given feasible solution $G$ is equal to:

$$\underbrace{\sum_{t \in T} |\overline{G}_t|}_{\text{add-moves}} + \underbrace{\sum_{\substack{t \in T \\ |G_t| \geq 2}} |G_t|}_{\text{remove-moves}} + \underbrace{\sum_{t \in T} \left( |G_t| \sum_{k=1}^{\min\{2, |\overline{G}_t|\}} \binom{|\overline{G}_t|}{k} \right)}_{\text{swap-moves}},$$

which is of order $O(|T| \cdot N^3)$ meaning that asymptotic complexity is dominated by the swap moves.

### 4 Data processing

#### 4.1 Filtering

**SARS-CoV-2.** We applied different quality-based filtering steps using the metadata provided with the genomes obtained from GISAID [4] on November 16th, 2025. For both the USA-based and China-based datasets, we filtered genomes based on the following criteria:

- *location* – USA or China
- *collection date* – between January 1st, 2022 and June 30th, 2024 (including endpoints)
- *ambiguous nucleotides* – $\leq$ 0.1% N-count and $\leq$ 1% non-ACGT-count
- *genome length* – $\geq$ 29,000bp
- *host* – Human host only
- *completeness* – this is a binary GISAID feature which we require to be "true"
- *lineage* we required the lineage to not be empty or equivalently undetermined

**IAV** We downloaded all human-host H1N1pdm09 and H3N2 genomes containing at least the complete HA and NA segments, with collection dates between January 1st, 2024 and March 31st, 2025 on November 20th, 2025. The filtering step for IAV was identical to that of SARS-CoV-2, with the exception that collection dates differed, and for IAV we excluded length-based and location-based filtering.

#### 4.2 Downsampling

**SARS-CoV-2.** As the number of USA-based SARS-CoV-2 genomes exceeded 100,000, we decided to downsample these using Algorithm 1 to retain at most $k = 3{,}500$ genomes per month with a similar lineage distribution as all genomes in the corresponding month. The implementation details for the downsampling procedure as well as additional steps can be found on our github https://github.com/JaspervB-tud/ReSeT under the manuscript folder.

**IAV.** For IAV, we used the same downsampling scheme, but instead of retaining 3,500 genomes per month, we reduced this to 2,000 per month.

**Algorithm 1:** Temporal downsampling

**Input:** Filtered genomes $G$ with collection dates and taxon assignments; monthly cap $k$; random seed
**Output:** Downsampled genome set $D$

$D \leftarrow \emptyset$;
$\mathcal{M} \leftarrow \textsc{GroupByMonth}(G)$;
**foreach** *month* $m \in \mathcal{M}$ **do**
  $\mathcal{T} \leftarrow \textsc{GroupByTaxon}(\mathcal{M}[m])$;
  $k_m \leftarrow \min\big(k,\ |\mathcal{M}[m]|\big)$;
  `// assign at least one genome per taxon`
  $a_t \leftarrow 1\ \forall, t \in \mathcal{T}$;
  `// determine remaining capacity and proportionally distribute over taxa`
  $\mathrm{cap}_t \leftarrow |\mathcal{T}[t]| - a_t\ \ \forall t$ ; $r \leftarrow k_m - \sum_t a_t$;
  $T \leftarrow \sum_t \mathrm{cap}_t$;
  **if** $r > 0$ ***and*** $T > 0$ **then**
    `// calculate proportions per taxon`
    $p_t \leftarrow r \cdot \mathrm{cap}_t \,/\, T\ \ \forall t$;
    `// determine number of genomes to select per taxon`
    $a_t \leftarrow a_t + \min\big(\mathrm{cap}_t,\ \lfloor p_t \rfloor\big)\ \ \forall t$;
    `// for leftover capacity, distribute over taxa based on largest fractional proportion`
    Sort taxa by $p_t - \lfloor p_t \rfloor$ descending;
    **foreach** $t$ ***while*** $\sum_t a_t < k_m$ **do**
      $a_t \leftarrow a_t + 1$;
    **end**
  **end**
  `// Randomly select genomes for every taxon`
  **foreach** $t \in \mathcal{T}$ ***with*** $a_t > 0$ **do**
    $D \leftarrow D \cup \textsc{RandomSample}(\mathcal{T}[t],\ a_t)$;
  **end**
**end**
**return** $D$

#### 4.3 Splitting the data into Reference and Simulation genomes

**SARS-CoV-2.** For SARS-CoV-2, we used December 31st, 2023 as the temporal cut-off to split reference and simulation genomes. Genomes with a collection date up until (including) December 31st, 2023 are considered reference genome candidates, and genomes with a collection date after are considered potential simulation genomes. For the simulation genomes, we exclude any genome with a lineage that does not occur in the reference data, as these will always be estimating at an abundance of 0.

**IAV.** For IAV we instead used December 31st, 2024 as the temporal cut-off, meaning that genomes with a collection date before 2025 are candidate reference genomes, and genomes with a later collection date are potential simulation genomes. As for SARS-CoV-2, we only retain genomes with a clade that occurs in the reference genomes.

#### 4.4 Simulation genome selection

##### 4.4.1 Distance calculations.

For both SARS-CoV-2 and IAV we wanted to determine distances in a way that did not rely on MASH [8] and sourmash [1], which are used for reference genome selection. To this end, we ran MAFFT v7.525 [3] to obtain multiple sequence alignments (MSAs) of the potential simulation genomes. For IAV, this was done per segment. Distances between genomes were then calculated from the MSA by parsing per column, considering only columns where at least one genome did not have a gap, and neither sequence had an ’N’. The final distance was given by dividing the number of mismatches by the total number of considered columns.

##### 4.4.2 Difficulty-stratified experiments

Due to the structural differences between SARS-CoV-2 and IAV, our approach differs between the viruses. However, in both cases we first eliminate duplicate genomes by building a graph where we draw an edge between a pair of sequences if they have a distance of 0. Afterwards, we perform a depth-first search to find connected components in this graph, and for every such component we only retain the genome with the lowest index (indices are determined by sorted sequence IDs). In what follows, we will describe the specific approaches for selecting genomes per virus.

**SARS-CoV-2.** For SARS-CoV-2, which has many lineages that are closely related, we utilize a clustering strategy to capture local similarity structures using HDBSCAN v0.8.39 [7] which creates density-based clusters consisting of at least 5 genomes per cluster (see Algorithm 2). Within each cluster, we determine lineage medoids after which we check for every constituent genome whether it is close to another lineage’s medoid than its own. If this is the case, the genome is considered *difficult* (since it is taxonomically ambiguous with respect to the cluster), and its lineage as well as the lineage of the medoid it is closer to is stored as an edge in a lineage-oriented conflict graph, indicating that this lineage pair is mutually confusable.

After assigning difficulty labels and finalizing the conflict graph, we want to find a maximum-weight independent set of non-isolated lineages in the graph (Algorithm 3). This choice is motivated by the fact that abundance estimation errors for lineages which are mutually confusable can be invisible as reads belonging to them are likely to be swapped between them. From the maximum independent set we retain up to $k_L = 10$ lineages (based of their respective difficulty), and from the lineages with degree zero in the conflict graph (i.e. lineages without confusable taxa) we retain the 10 largest lineages. For the 10 hard lineages we sort genomes first by whether they are labeled as difficult and second by the fraction of genomes of the same cluster with a different lineage that are closer to it, than it is to its own medoid within its cluster. Hard genomes are obtained by taking the top-$k_\ell = 5$ genomes of this sorted list, which are labeled as difficult (up to 5 if fewer are available). The bottom 5 genomes, regardless of difficulty status, comprise the *Medium* genomes for this lineage. Finally, for the *Easy* genomes, we take the 5 genomes with largest minimum distance (not restricted to clustering) to any genome of another lineage.

**Algorithm 2:** Difficulty-based selection of SARS-CoV-2 simulation genomes

**Input:** Distance matrix $\mathbf{D}$ over simulation genomes; lineage assignments; max. genomes per lineage $k_\ell$; max. lineages $k_L$

**Output:** Hard, medium, and easy genome sets $S_H$, $S_M$, $S_E$

Remove exact duplicates via connected components with $d = 0$;
Cluster genomes using HDBSCAN (MIN_CLUSTER_SIZE=5) on $\mathbf{D}$; assign unclustered points to singleton clusters;
Initialize conflict graph $\mathcal{G}$ with one node per lineage;
**foreach** *cluster $C$* **do**
  **foreach** *lineage $\ell$ present in $C$* **do**
    $\mu_\ell^C \leftarrow \arg\min_{i \in C_\ell} \sum_{j \in C_\ell} D_{ij}$ `// medoid of` $\ell$ `in` $C$
  **end**
  **foreach** *genome $g \in C$ with true lineage $\ell$* **do**
    Record $d_g \leftarrow D(g, \mu_\ell^C)$;
    **foreach** *other lineage $\ell' \neq \ell$ in $C$* **do**
      **if** $D(g, \mu_{\ell'}^C) < d_g$ **then**
        Mark $g$ as *difficult*;
        Add edge $(\ell, \ell')$ to $\mathcal{G}$;
      **end**
    **end**
  **end**
**end**
$\mathcal{H}, \mathcal{E} \leftarrow$ GreedyMWIS($\mathcal{G}$) `// see Algorithm 3`
Retain the top-$k_L$ lineages from $\mathcal{H}$ and $\mathcal{E}$ by number of difficult and total genomes, respectively;
**foreach** *lineage $\ell \in \mathcal{H}$* **do**
  **foreach** *genome $g$ in lineage $\ell$* **do**
    score($g$) $\leftarrow$ fraction of within-cluster neighbours from foreign lineages that are closer to $g$ than $d_g$;
  **end**
  Sort genomes by (is difficult, score) descending;
  $S_H[\ell] \leftarrow$ top $\min(k_\ell, |\{\text{difficult in } \ell\}|)$ genomes;
  $S_M[\ell] \leftarrow$ bottom $k_\ell$ genomes by score;
**end**
**foreach** *lineage $\ell \in \mathcal{E}$* **do**
  $S_E[\ell] \leftarrow k_\ell$ genomes with largest minimum distance to any foreign genome;
**end**
**return** $S_H$, $S_M$, $S_E$

**Algorithm 3:** GreedyMWIS: greedy maximum-weight independent set on conflict graph

**Input:** Conflict graph $\mathcal{G}$; number of difficult genomes per lineage $n_\ell^d$
**Output:** Hard lineage set $\mathcal{H}$; easy lineage set $\mathcal{E}$
`// assign weights based on number of difficult genomes per lineage`
**foreach** *lineage* $\ell \in \mathcal{G}$ **do**

$$w_\ell \leftarrow \begin{cases} 0 & \text{if } n_\ell^d = 0 \\ 0.5 & \text{if } 1 \leq n_\ell^d \leq 5 \\ 1 & \text{if } 5 < n_\ell^d \leq 15 \\ 3 & \text{if } n_\ell^d > 15 \end{cases};$$

**end**
$\mathcal{E} \leftarrow \{\ell \in \mathcal{G} : \deg(\ell) = 0\}$;
$\mathcal{G} \leftarrow \mathcal{G} \setminus \mathcal{E}$;
$\mathcal{H} \leftarrow \emptyset$;
**while** $\mathcal{G}$ *has nodes* **do**
    `// select lineage with highest weight normalized by degree`
    $\ell^* \leftarrow \arg\max_\ell \; w_\ell \,/\, (\deg(\ell) + 1)$;
    **if** $w_{\ell^*} > 0$ **then**
        $\mathcal{H} \leftarrow \mathcal{H} \cup \{\ell^*\}$;
    **end**
    Remove $\ell^*$ and all its neighbours from $\mathcal{G}$;
**end**
**return** $\mathcal{H}$, $\mathcal{E}$

**IAV.** The methodology defined above was infeasible for IAV due to better clade separation, and a low number of clades ($< 10$). Therefore, we utilized a simpler approach: we calculate the medoid genome of every clade, and for every genome calculate the distance to its clade's medoid ($d_{\text{intra}}$), and the closest other clade's medoid ($d_{\text{inter}}$). Based on these distances, we calculate $d_{\text{intra}} - d_{\text{inter}}$, which acts as a proxy for the difficulty score of every sequence. Sorting sequences by this scoring, we define Hard sequences as having a non-negative score (i.e. $d_{\text{intra}} \geq d_{\text{inter}}$), and Easy sequences as having a negative score. Then, for every clade we pick the top-5 sequences with a positive score as *Hard* sequences, and the bottom-5 as *Easy* sequences.

###### 4.4.3 Dirichlet-distributed experiments

For the Dirichlet-distributed experiments, we generated samples for three symmetric Dirichlet distributions (concentration $\alpha \in \{0.1, 1.0, 10.0\}$). We create 10 compositions using 20 randomly selected SARS-CoV-2 lineages, or 5 randomly selected IAV clades. These compositions will be kept constant across $\alpha$ values, allowing us to isolate the effect of different lineage abundance distributions. For every taxon (lineage/clade), we select up to 10 genomes (based on availability) by first taking the medoid (using the distances obtained from the MSA), and then iteratively adding the genome most distant from all previously selected genomes within the taxon.

#### 4.5 Parameter tuning

##### 4.5.1 Splitting the Reference genomes into Train and Test genomes

Since the final simulation genomes should not be used to perform parameter tuning, we instead create sets of *Train* and *Test* genomes from the Reference genome set for every dataset. In this, we mimic the temporal split used to partition the Reference and Simulation genomes, and we downsample the number of genomes to keep computations feasible.

**SARS-CoV-2.** For SARS-CoV-2, we restrict Train genomes to have a collection date between January 1st, 2022 and December 31st, 2022, whereas Test genomes have a collection date between January 1st, 2023 and June 30th, 2023. Moreover, to keep computations fast we randomly subsample up to 200 train genomes and 100 test genomes per lineage that occurs in the train set.

**IAV.** The treatment for IAV is along the same lines as that of SARS-CoV-2: we restrict Train genomes to have a collection date between January 1st, 2024 and September 31st, 2024 whereas Test genomes have a collection date between October 1st, 2024 and December 31st, 2024. As there are fewer than 10 clades, we downscale to at most 1,000 train genomes and 500 test genomes per clade.

##### 4.5.2 Parameter grids

For all three datasets, we use the same baseline methods: VSEARCH v2.28.1 [9], complete-linkage hierarchical clustering using SciPy v1.5.0 https://scipy.org, medoid-based selection and no selection (i.e. including all reference genomes). The latter of these selections does not have any parameters, and was therefore excluded from the parameter tuning. For all other baseline methods, we consider a grid of tuning parameters and for each method and parameter combination we compute the corresponding reference genomes. Specifically, for distance estimation methods we consider MASH v2.3 [8] using two different sketch sizes per virus (5,000 and 10,000 for SARS-CoV-2, and 500 and 1,000 for IAV) and a fixed $k$-mer size of 31, and we consider sourmash using version 0.9.14 of the branchwater plugin [1] with the same fixed $k$-mer size of 31 and scale values of 6 and 3. Sourmash operates on a slightly different basis than MASH, and determines both Jaccard and cosine similarities (that can be turned into dissimilarity score by subtracting them from 1), which we both consider throughout parameter tuning.

**Hierarchical clustering.** For hierarchical clustering specifically, we consider both *static* and *dynamic* similarity thresholds for every distance estimation method included. Static similarity thresholds are fixed a priori and constant for every taxon. Dynamic thresholds, instead, parse the distance matrix on a per-taxon basis and determines a threshold for every individual taxon by calculating percentiles of all non-zero distances. For SARS-CoV-2, we consider static similarity thresholds of 99%, 99.9%, 99.99% and 99.999% and dynamic thresholds based on the 1st, 25th, 50th, 75th and 90th percentile. Due to lower overall similarities, we instead use static thresholds of 95%, 97%, 99% for IAV, while using the same percentiles for dynamic thresholds. After clustering, the medoid of every cluster is selected as a reference genome.

**Medoid.** For medoid selection we simply consider the medoid sequence for every taxon, based on the distance/similarity estimation techniques described above.

**VSEARCH.** Similar to hierarchical clustering, we use both static and dynamic similarity thresholds for VSEARCH. However, as VSEARCH internally determines distances based on exact alignment, we use dynamic thresholds obtained for hierarchical clustering (computed with MASH and sourmash), resulting in one selection for every dynamic threshold en distance estimation method-parameter combination. For both viruses, we use the same thresholds as for hierarchical clustering.

**ReSeT.** ReSeT has a selection cost $c$, and a scale parameter $\lambda$ which assigns weight to the inter-taxon penalization in our selection model. For parameter tuning we consider different orders of magnitude for both parameters (in combination with every distance estimation method), so that $c \in \{10^0, 10^{-1}, 10^{-2}, 10^{-3}, 10^{-4}, 10^{-5}, 10^{-6}, 0\}$ and $\lambda \in \{10^0, 10^{-1}, 10^{-2}, 10^{-3}, 10^{-4}, 10^{-5}, 0\}$. When $\lambda$ is set to 0, our model ignores inter-taxon penalties entirely, and the model simplifies to the standard facility location problem model per taxon that is also utilized by PARNAS [6] and Repset [5].

##### 4.5.3 Evaluation and weighing

For every method and parameter combination, we compute a reference set by running the respective selection methods and aggregating genomes over all taxa. Based on these reference sets we profile the simulated difficulty-stratified tuning samples as described in the main text to obtain abundance accuracies and F1-scores. For both metrics, we take the median score per difficulty level and compute a weighted average score across difficulties. For SARS-CoV-2, we apply weights $w_{\text{Easy}} = 0.2, w_{\text{Medium}} = 0.5, w_{\text{Hard}} = 0.3$ and average over the USA-based and China-based datasets (equal weight). For IAV, we use $w_{\text{Easy}} = 0.4, w_{\text{Hard}} = 0.6$. We then rank all method-parameter combinations by their weighted abundance accuracy (ties broken by weighted F1-score) and by their weighted F1-score (ties broken by weighted abundance accuracy). Additionally, we apply a minimax-rank criterion where, for every parameterization, we take its worst rank across both weighted metrics and select the parameterization with the best worst-case rank. This results in up to three selections per method per virus.

#### 4.6 Parameter tuning outcomes

##### 4.6.1 SARS-CoV-2

**Supplementary Table S1:** All medoid parameterizations for SARS-CoV-2, ordered by minimax rank. Selected parameterizations are shown in bold, highlighting the metric on which they were selected.

| Method | $s$ | $r_{\mathrm{mm}}$ | Weighted Abund. | $r_{\mathrm{Abund.}}$ | Weighted F1 | $r_{\mathrm{F1}}$ |
|---|---|---|---|---|---|---|
| **MASH** | **10,000** | **1** | **0.561** | **1** | **0.307** | **1** |
| Sourmash Jaccard | 3 | 3 | 0.543 | 3 | 0.302 | 3 |
| Sourmash Cosine | 3 | 4 | 0.538 | 4 | 0.300 | 4 |
| MASH | 5,000 | 5 | 0.538 | 5 | 0.306 | 2 |
| Sourmash Jaccard | 6 | 5 | 0.553 | 2 | 0.282 | 5 |
| Sourmash Cosine | 6 | 6 | 0.520 | 6 | 0.281 | 6 |

**Supplementary Table S2:** Top-10 hierarchical clustering parameterizations for SARS-CoV-2, ordered by minimax rank. Selected parameterizations are shown in bold, highlighting the metric on which they were selected.

| Distance | $s$ | $t$ | $r_{\mathrm{mm}}$ | Weighted Abund. | $r_{\mathrm{Abund.}}$ | Weighted F1 | $r_{\mathrm{F1}}$ |
|---|---|---|---|---|---|---|---|
| **MASH** | **10,000** | **0.99** | **1** | **0.561** | **1** | **0.307** | **1** |
| MASH | 10,000 | 0.999 | 3 | 0.555 | 2 | 0.303 | 3 |
| MASH | 5,000 | 0.999 | 5 | 0.541 | 5 | 0.303 | 4 |
| MASH | 5,000 | 0.99 | 7 | 0.538 | 7 | 0.306 | 2 |
| Sourmash Jaccard | 3 | 90 | 7 | 0.549 | 3 | 0.238 | 7 |
| Sourmash Cosine | 3 | 90 | 8 | 0.543 | 4 | 0.235 | 8 |
| Sourmash Cosine | 6 | 75 | 9 | 0.540 | 6 | 0.233 | 9 |
| MASH | 5,000 | 90 | 10 | 0.537 | 9 | 0.233 | 10 |
| Sourmash Jaccard | 6 | 75 | 12 | 0.537 | 8 | 0.230 | 12 |
| MASH | 10,000 | 50 | 22 | 0.536 | 10 | 0.206 | 22 |

**Supplementary Table S3:** Top-10 VSEARCH parameterizations for SARS-CoV-2, ordered by minimax rank. 14 additional Sourmash parameterizations (all remaining combinations of Jaccard/Cosine, $s \in \{3, 6\}$, $t \in \{25, 50, 75, 90\}$) produced identical selections to the selected F1- and minimax-based selections, and are collapsed into a single entry in the table. Selected parameterizations are shown in bold, highlighting the metric on which they were selected.

| Distance | $s$ | $t$ | $r_{\mathrm{mm}}$ | Weighted Abund. | $r_{\mathrm{Abund.}}$ | Weighted F1 | $r_{\mathrm{F1}}$ |
|---|---|---|---|---|---|---|---|
| **Sourmash Cosine**[a] | **3** | **1** | 13 | 0.466 | 13 | **0.270** | **1** |
| **Sourmash Jaccard**[b] | **6** | **25** | **13** | 0.466 | 13 | 0.270 | 1 |
| Sourmash | – | – | 13 | 0.466 | 13 | 0.270 | 1 |
| Sourmash Jaccard | 6 | 1 | 17 | 0.522 | 4 | 0.269 | 17 |
| Sourmash Jaccard | 3 | 1 | 20 | 0.521 | 6 | 0.262 | 20 |
| Default | – | 0.999 | 22 | 0.512 | 7 | 0.261 | 22 |
| MASH | 10,000 | 90 | 23 | 0.526 | 3 | 0.241 | 23 |
| **MASH** | **5,000** | **90** | 24 | **0.546** | **1** | 0.239 | 24 |
| MASH | 10,000 | 75 | 25 | 0.521 | 5 | 0.230 | 25 |
| MASH | 5,000 | 75 | 26 | 0.527 | 2 | 0.225 | 26 |

[a] Selected parameterization for F1-score.
[b] Selected parameterization for minimax-rank.

**Supplementary Table S4:** Top-8 ReSeT parameterizations for SARS-CoV-2, ordered by minimax rank. As the best abundance accuracy configuration had a weighted $F1$ rank larger than 100, we only include the top 8 ordered by minimax rank and skip to the corresponding best abundance accuracy configuration. Selected parameterizations are shown in bold, highlighting the metric on which they were selected.

| Distance | $s$ | $\lambda$ | $c$ | $r_{\text{mm}}$ | Weighted Abund. | $r_{\text{Abund.}}$ | Weighted F1 | $r_{\text{F1}}$ |
|---|---|---|---|---|---|---|---|---|
| **Mash** | **5,000** | **0** | **1** | **12** | 0.555 | 12 | 0.308 | 3 |
| **Mash** | **10,000** | $\mathbf{10^{-5}}$ | $\mathbf{10^{-1}}$ | 16 | 0.554 | 16 | **0.308** | **1** |
| Mash | 10,000 | $10^{-4}$ | 1 | 17 | 0.554 | 17 | 0.306 | 12 |
| Mash | 10,000 | 0 | $10^{-1}$ | 21 | 0.554 | 20 | 0.305 | 21 |
| Mash | 10,000 | 0 | 1 | 22 | 0.554 | 21 | 0.305 | 22 |
| Mash | 5,000 | 0 | $10^{-2}$ | 22 | 0.554 | 22 | 0.306 | 9 |
| Mash | 5,000 | $10^{-5}$ | 1 | 24 | 0.554 | 19 | 0.304 | 24 |
| Mash | 10,000 | $10^{-3}$ | $10^{-1}$ | 24 | 0.554 | 24 | 0.308 | 4 |
| Mash | 10,000 | $10^{-3}$ | 1 | 25 | 0.554 | 25 | 0.308 | 5 |
| | | | | ... | | | | |
| **Sourmash Cosine** | **3** | $\mathbf{10^{-5}}$ | $\mathbf{10^{-1}}$ | > 100 | **0.585** | **1** | 0.242 | > 100 |

##### 4.6.2 IAV

**Supplementary Table S5:** All medoid parameterizations for IAV, ordered by minimax rank. Selected parameterizations are shown in bold, highlighting the metric on which they were selected.

| Method | $s$ | $r_{\text{mm}}$ | Weighted Abund. | $r_{\text{Abund.}}$ | Weighted F1 | $r_{\text{F1}}$ |
|---|---|---|---|---|---|---|
| **MASH** | **500** | **1** | **0.666** | **1** | **0.897** | **1** |
| Sourmash Jaccard | 3 | 2 | 0.617 | 2 | 0.897 | 1 |
| Sourmash Jaccard | 6 | 3 | 0.615 | 3 | 0.897 | 1 |
| Sourmash Cosine | 6 | 3 | 0.615 | 3 | 0.897 | 1 |
| Sourmash Cosine | 3 | 5 | 0.603 | 5 | 0.897 | 1 |
| MASH | 1,000 | 6 | 0.602 | 6 | 0.897 | 1 |

**Supplementary Table S6:** Top-10 hierarchical clustering parameterizations for IAV, ordered by minimax rank. Selected parameterizations are shown in bold, highlighting the metric on which they were selected.

| Distance | $s$ | $t$ | $r_{\text{mm}}$ | Weighted Abund. | $r_{\text{Abund.}}$ | Weighted F1 | $r_{\text{F1}}$ |
|---|---|---|---|---|---|---|---|
| **MASH** | **1,000** | **90** | **1** | **0.718** | **1** | **0.943** | **1** |
| MASH | 500 | 90 | 2 | 0.694 | 2 | 0.943 | 1 |
| Sourmash Cosine | 3 | 90 | 5 | 0.659 | 5 | 0.943 | 1 |
| Sourmash Jaccard | 3 | 90 | 6 | 0.659 | 6 | 0.943 | 1 |
| MASH | 1,000 | 0.99 | 7 | 0.653 | 7 | 0.943 | 1 |
| Sourmash Jaccard | 3 | 25 | 8 | 0.649 | 8 | 0.943 | 1 |
| Sourmash Cosine | 3 | 25 | 9 | 0.645 | 9 | 0.943 | 1 |
| MASH | 500 | 25 | 10 | 0.640 | 10 | 0.943 | 1 |
| MASH | 1,000 | 25 | 11 | 0.640 | 11 | 0.943 | 1 |
| MASH | 500 | 50 | 12 | 0.638 | 12 | 0.943 | 1 |

**Supplementary Table S7:** Top-10 VSEARCH parameterizations, ordered by minimax rank. 20 additional parameterizations produced identical scores and are collapsed into a single entry: 16 Sourmash (all Jaccard/Cosine combinations, $s \in \{3, 6\}$, $t \in \{25, 50, 75, 90\}$), Sourmash Cosine ($s \in \{3, 6\}$, $t = 1$), and static thresholds ($t \in \{0.95, 0.97\}$). Selected parameterizations are shown in bold, highlighting the metric on which they were selected.

| Distance | $s$ | $t$ | $r_{\text{mm}}$ | Weighted Abund. | $r_{\text{Abund.}}$ | Weighted F1 | $r_{\text{F1}}$ |
|---|---|---|---|---|---|---|---|
| **Default**[a,b] | – | **0.99** | **2** | 0.800 | 2 | **0.943** | **1** |
| MASH | 500 | 50 | 3 | 0.733 | 3 | 0.943 | 1 |
| **MASH** | **1,000** | **75** | 5 | **0.802** | **1** | 0.897 | 5 |
| MASH | 500 | 90 | 5 | 0.725 | 4 | 0.897 | 5 |
| MASH | 1,000 | 90 | 5 | 0.719 | 5 | 0.897 | 5 |
| MASH | 1,000 | 50 | 7 | 0.704 | 7 | 0.943 | 1 |
| Various[c] | – | – | 10 | 0.645 | 10 | 0.897 | 5 |
| MASH | 500 | 1 | 30 | 0.646 | 8 | 0.888 | 30 |
| MASH | 1,000 | 25 | 31 | 0.640 | 31 | 0.943 | 1 |
| MASH | 500 | 25 | 32 | 0.632 | 32 | 0.888 | 30 |

[a] Selected parameterization for F1-score.
[b] Selected parameterization for minimax rank.
[c] 20 parameterizations with identical scores: see caption for details.

**Supplementary Table S8:** Top-8 ReSeT parameterizations for IAV, ordered by minimax rank. As the best abundance accuracy configuration had a weighted $F1$ rank larger than 45, we only include the top 8 ordered by minimax rank and skip to the corresponding best abundance accuracy configuration. Selected parameterizations are shown in bold, highlighting the metric on which they were selected.

| Distance | $s$ | $\lambda$ | $c$ | $r_{\text{mm}}$ | Weighted Abund. | $r_{\text{Abund.}}$ | Weighted F1 | $r_{\text{F1}}$ |
|---|---|---|---|---|---|---|---|---|
| **MASH**[a,b] | **500** | **0** | $\mathbf{10^{-2}}$ | **5** | 0.648 | 5 | **0.943** | **1** |
| MASH | 1,000 | 0 | $10^{-2}$ | 6 | 0.647 | 6 | 0.943 | 1 |
| MASH | 500 | $10^{-5}$ | $10^{-2}$ | 7 | 0.647 | 7 | 0.943 | 1 |
| MASH | 500 | $10^{-4}$ | $10^{-2}$ | 7 | 0.647 | 7 | 0.943 | 1 |
| MASH | 500 | $10^{-3}$ | $10^{-2}$ | 9 | 0.647 | 9 | 0.943 | 1 |
| MASH | 1,000 | $10^{-3}$ | $10^{-2}$ | 10 | 0.646 | 10 | 0.943 | 1 |
| MASH | 1,000 | $10^{-5}$ | $10^{-2}$ | 11 | 0.646 | 11 | 0.943 | 1 |
| MASH | 1,000 | $10^{-4}$ | $10^{-2}$ | 11 | 0.646 | 11 | 0.943 | 1 |
| | | | | | ... | | | |
| **MASH**[c] | **500** | **0** | **1** | >45 | **0.666** | **1** | 0.897 | >45 |

[a] Selected parameterization for F1-score.
[b] Selected parameterization for minimax rank.
[c] Selected parameterization for abundance accuracy (L1).

#### 4.7 Effect sizes for difficulty-based experiments

**Supplementary Table S9:** Statistical comparison of ReSeT configurations against the best non-ReSeT baseline per dataset on difficulty samples: abundance accuracy (paired Wilcoxon signed-rank test, two-sided; $n = 20$ per difficulty). $\Delta$ denotes the difference in median abundance accuracy between ReSeT and the baseline (positive = ReSeT outperforms). $p$: raw two-sided Wilcoxon $p$-value. $q$: Benjamini–Hochberg-adjusted $p$-value within the per-difficulty family for each dataset (BH family size: 18 for SARS-CoV-2, 8 for IAV). $p < 10^{-5}$ is the minimum achievable $p$-value at $n = 20$. Significance markers: $^{*}q < 0.05$, $^{**}q < 0.01$, $^{***}q < 0.001$. Cells in **bold** indicate significant improvements over the baseline. Baselines: Medoid/Hierarchical clustering (MASH, $s = 10{,}000$) for SARS-CoV-2; VSEARCH (threshold 0.99) for IAV.

| | Easy | | | Medium | | | Hard | | |
|---|---|---|---|---|---|---|---|---|---|
| ReSeT config | $\Delta$ Abund. | $p$ | $q$ | $\Delta$ Abund. | $p$ | $q$ | $\Delta$ Abund. | $p$ | $q$ |
| *SARS-CoV-2 (USA)* | | | | | | | | | |
| ReSeT (F1) | +**0.012**$^{***}$ | $1.3\times10^{-5}$ | $3.0\times10^{-5}$ | +**0.088**$^{***}$ | $<10^{-5}$ | $<10^{-5}$ | +**0.088**$^{***}$ | $<10^{-5}$ | $<10^{-5}$ |
| ReSeT (Abund.) | −0.176$^{***}$ | $<10^{-5}$ | $<10^{-5}$ | +0.004 | $4.3\times10^{-1}$ | $4.8\times10^{-1}$ | +0.007 | $8.4\times10^{-1}$ | $8.4\times10^{-1}$ |
| ReSeT (Rank) | −0.034$^{***}$ | $<10^{-5}$ | $<10^{-5}$ | +**0.062**$^{***}$ | $<10^{-5}$ | $<10^{-5}$ | +**0.093**$^{***}$ | $<10^{-5}$ | $<10^{-5}$ |
| *SARS-CoV-2 (China)* | | | | | | | | | |
| ReSeT (F1) | +0.001 | $4.0\times10^{-2}$ | $9.0\times10^{-2}$ | −0.002 | $3.7\times10^{-1}$ | $4.7\times10^{-1}$ | −0.002 | $4.8\times10^{-2}$ | $9.7\times10^{-2}$ |
| ReSeT (Abund.) | −0.075$^{***}$ | $<10^{-5}$ | $1.1\times10^{-5}$ | +**0.088**$^{***}$ | $<10^{-5}$ | $1.1\times10^{-5}$ | +**0.066**$^{***}$ | $<10^{-5}$ | $1.1\times10^{-5}$ |
| ReSeT (Rank) | +0.001 | $9.0\times10^{-2}$ | $1.6\times10^{-1}$ | +0.001 | $3.9\times10^{-1}$ | $4.7\times10^{-1}$ | +0.015 | $1.1\times10^{-1}$ | $1.6\times10^{-1}$ |
| *IAV* | | | | | | | | | |
| ReSeT (F1/Rank) | −0.293$^{***}$ | $<10^{-5}$ | $<10^{-5}$ | — | — | — | −0.041$^{***}$ | $<10^{-5}$ | $<10^{-5}$ |
| ReSeT (Abund.) | −0.426$^{***}$ | $<10^{-5}$ | $<10^{-5}$ | — | — | — | −0.102$^{***}$ | $<10^{-5}$ | $<10^{-5}$ |

**Supplementary Table S10:** Statistical comparison of ReSeT configurations against the best non-ReSeT baseline per dataset on difficulty samples: F1-score (paired Wilcoxon signed-rank test, two-sided; $n = 20$ per difficulty). All conventions as in Table S9.

| | Easy | | | Medium | | | Hard | | |
|---|---|---|---|---|---|---|---|---|---|
| ReSeT config | $\Delta$ F1-score | $p$ | $q$ | $\Delta$ F1-score | $p$ | $q$ | $\Delta$ F1-score | $p$ | $q$ |
| *SARS-CoV-2 (USA)* | | | | | | | | | |
| ReSeT (F1) | −0.004 | $9.3\times10^{-2}$ | $1.2\times10^{-1}$ | +**0.022**$^{***}$ | $<10^{-5}$ | $<10^{-5}$ | +**0.019**$^{***}$ | $4.8\times10^{-4}$ | $7.2\times10^{-4}$ |
| ReSeT (Abund.) | −0.083$^{***}$ | $8.8\times10^{-5}$ | $1.6\times10^{-4}$ | −0.026$^{***}$ | $1.4\times10^{-4}$ | $2.3\times10^{-4}$ | −0.041$^{***}$ | $8.8\times10^{-5}$ | $1.6\times10^{-4}$ |
| ReSeT (Rank) | −0.006$^{*}$ | $1.1\times10^{-2}$ | $1.5\times10^{-2}$ | −0.009 | $6.0\times10^{-1}$ | $6.4\times10^{-1}$ | +0.005 | $1.1\times10^{-1}$ | $1.3\times10^{-1}$ |
| *SARS-CoV-2 (China)* | | | | | | | | | |
| ReSeT (F1) | +0.009 | $1.1\times10^{-1}$ | $1.6\times10^{-1}$ | +0.000 | $8.6\times10^{-1}$ | $9.1\times10^{-1}$ | +0.000 | $1.1\times10^{-1}$ | $1.6\times10^{-1}$ |
| ReSeT (Abund.) | −0.081$^{***}$ | $8.8\times10^{-5}$ | $2.6\times10^{-4}$ | −0.114$^{***}$ | $8.8\times10^{-5}$ | $2.6\times10^{-4}$ | −0.096$^{***}$ | $8.8\times10^{-5}$ | $2.6\times10^{-4}$ |
| ReSeT (Rank) | +0.000 | $6.7\times10^{-1}$ | $7.5\times10^{-1}$ | +0.000 | $9.5\times10^{-1}$ | $9.5\times10^{-1}$ | +**0.011**$^{**}$ | $1.2\times10^{-3}$ | $3.2\times10^{-3}$ |
| *IAV* | | | | | | | | | |
| ReSeT (F1/Rank) | −0.091$^{***}$ | $1.3\times10^{-5}$ | $2.1\times10^{-5}$ | — | — | — | +**0.190**$^{***}$ | $3.6\times10^{-5}$ | $4.2\times10^{-5}$ |
| ReSeT (Abund.) | −0.231$^{***}$ | $3.7\times10^{-5}$ | $4.2\times10^{-5}$ | — | — | — | +**0.190**$^{***}$ | $4.8\times10^{-5}$ | $4.8\times10^{-5}$ |

#### 4.8 Rankings for per-alpha Dirichlet-distributed samples

**Supplementary Table S11:** Mean ranks for reference selection methods across Dirichlet-distributed compositions (10 compositions) per concentration parameter. Methods are ranked per composition (rank 1 = best). Best value per column in **bold**.

| | F1 | | | Abund. | | | Avg | | |
|---|---|---|---|---|---|---|---|---|---|
| Method | $\alpha = 0.1$ | 1.0 | 10.0 | $\alpha = 0.1$ | 1.0 | 10.0 | $\alpha = 0.1$ | 1.0 | 10.0 |
| *SARS-CoV-2 (USA)* | | | | | | | | | |
| All | 7.80 | 7.50 | 7.10 | 6.00 | 8.00 | 8.00 | 6.90 | 7.75 | 7.55 |
| Medoid/Hier. | 2.70 | 2.40 | **1.90** | 4.10 | 3.40 | 3.90 | 3.40 | 2.90 | 2.90 |
| VSEARCH (F1) | 3.60 | 5.10 | 5.30 | 4.40 | 4.10 | 3.70 | 4.00 | 4.60 | 4.50 |
| VSEARCH (Abund.) | 5.50 | 5.10 | 5.60 | 4.10 | 5.00 | 4.50 | 4.80 | 5.05 | 5.05 |
| VSEARCH (Rank) | 3.60 | 5.10 | 5.30 | 4.40 | 4.10 | 3.70 | 4.00 | 4.60 | 4.50 |
| ReSeT (F1) | **2.40** | **1.70** | **1.90** | 4.80 | **2.70** | **3.20** | 3.60 | **2.20** | **2.55** |
| ReSeT (Abund.) | 6.90 | 6.80 | 6.20 | 4.20 | 5.50 | 5.40 | 5.55 | 6.15 | 5.80 |
| ReSeT (Rank) | 3.50 | 2.30 | 2.70 | **4.00** | 3.20 | 3.60 | **3.75** | 2.75 | 3.15 |
| *SARS-CoV-2 (China)* | | | | | | | | | |
| All | 8.00 | 7.80 | 7.40 | 5.00 | 4.90 | 4.40 | 6.50 | 6.35 | 5.90 |
| Medoid/Hier. | 2.80 | 2.10 | **1.50** | 3.00 | 4.30 | 4.10 | 2.90 | 3.20 | 2.80 |
| VSEARCH (F1) | 6.00 | 6.50 | 6.80 | 7.20 | 6.00 | 7.30 | 6.60 | 6.25 | 7.05 |
| VSEARCH (Abund.) | 4.00 | 4.50 | 4.70 | 4.10 | 3.30 | **2.80** | 4.05 | 3.90 | 3.75 |
| VSEARCH (Rank) | 5.60 | 6.30 | 6.20 | 6.60 | 5.80 | 6.70 | 6.10 | 6.05 | 6.45 |
| ReSeT (F1) | 2.60 | 2.50 | 3.00 | 4.00 | 5.00 | 5.10 | 3.30 | 3.75 | 4.05 |
| ReSeT (Abund.) | 5.00 | 4.90 | 4.90 | 3.30 | 4.10 | 3.10 | 4.15 | 4.50 | 4.00 |
| ReSeT (Rank) | **2.00** | **1.40** | **1.50** | **2.80** | **2.60** | 2.50 | **2.40** | **2.00** | **2.00** |
| *IAV* | | | | | | | | | |
| All | 4.40 | 3.85 | 3.70 | 2.90 | 4.30 | 5.40 | 3.65 | 4.08 | 4.55 |
| Medoid | 3.65 | 5.00 | 5.20 | 3.90 | 4.00 | 2.40 | 3.77 | 4.50 | 3.80 |
| Hier. | 3.75 | 5.00 | 5.20 | 4.40 | 4.30 | 3.00 | 4.08 | 4.65 | 4.10 |
| VSEARCH (F1/Rank) | **2.40** | **1.70** | **1.50** | 3.80 | **2.00** | **1.10** | **3.10** | **1.85** | **1.30** |
| VSEARCH (Abund.) | 3.40 | 2.30 | 2.10 | 4.80 | 3.40 | 3.80 | 4.10 | 2.85 | 2.95 |
| ReSeT (F1/Rank) | 4.10 | 3.75 | 3.50 | **2.60** | 3.50 | 5.50 | 3.35 | 3.62 | 4.50 |
| ReSeT (Abund.) | 6.30 | 6.40 | 6.80 | 5.60 | 6.50 | 6.80 | 5.95 | 6.45 | 6.80 |

#### 4.9 Parameter tuning optimization landscapes for SARS-CoV-2

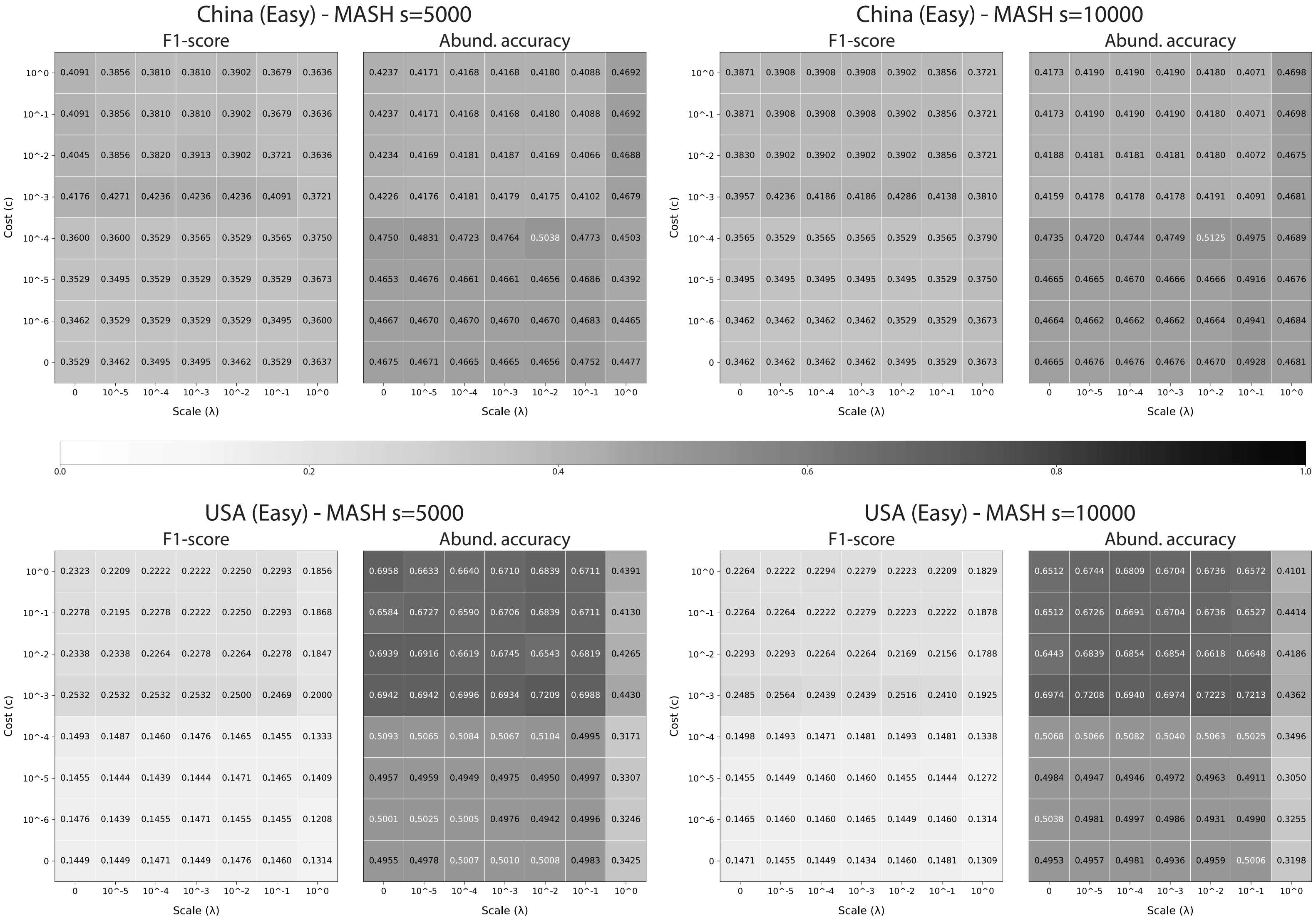

**Supplementary Figure S1:** Accuracy metrics for Easy samples across selection cost $c$ and scale parameter $\lambda$ during SARS-CoV-2 tuning experiments using MASH to estimate distances.

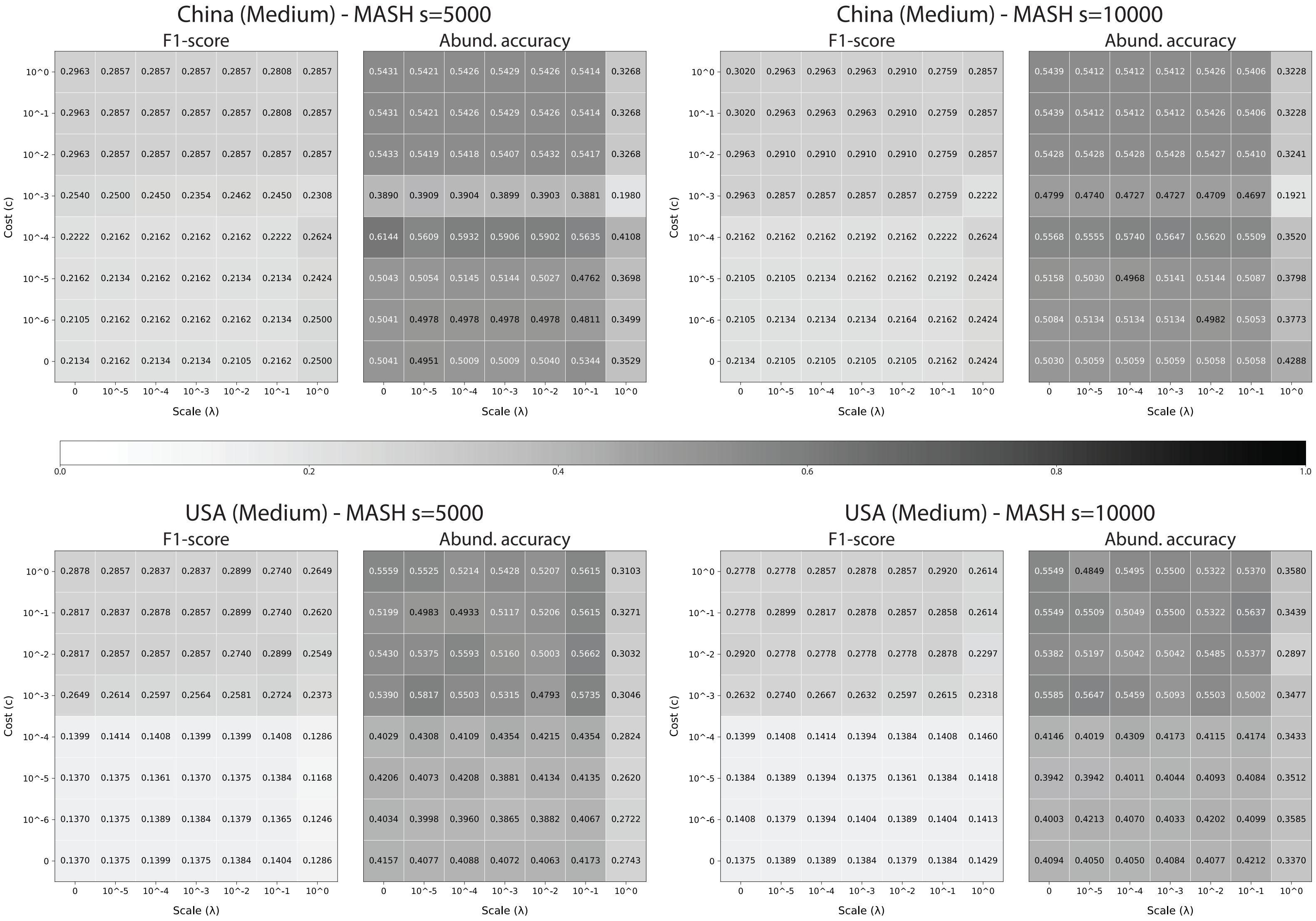

**Supplementary Figure S2:** Accuracy metrics for Medium samples across selection cost $c$ and scale parameter $\lambda$ during SARS-CoV-2 tuning experiments using MASH to estimate distances.

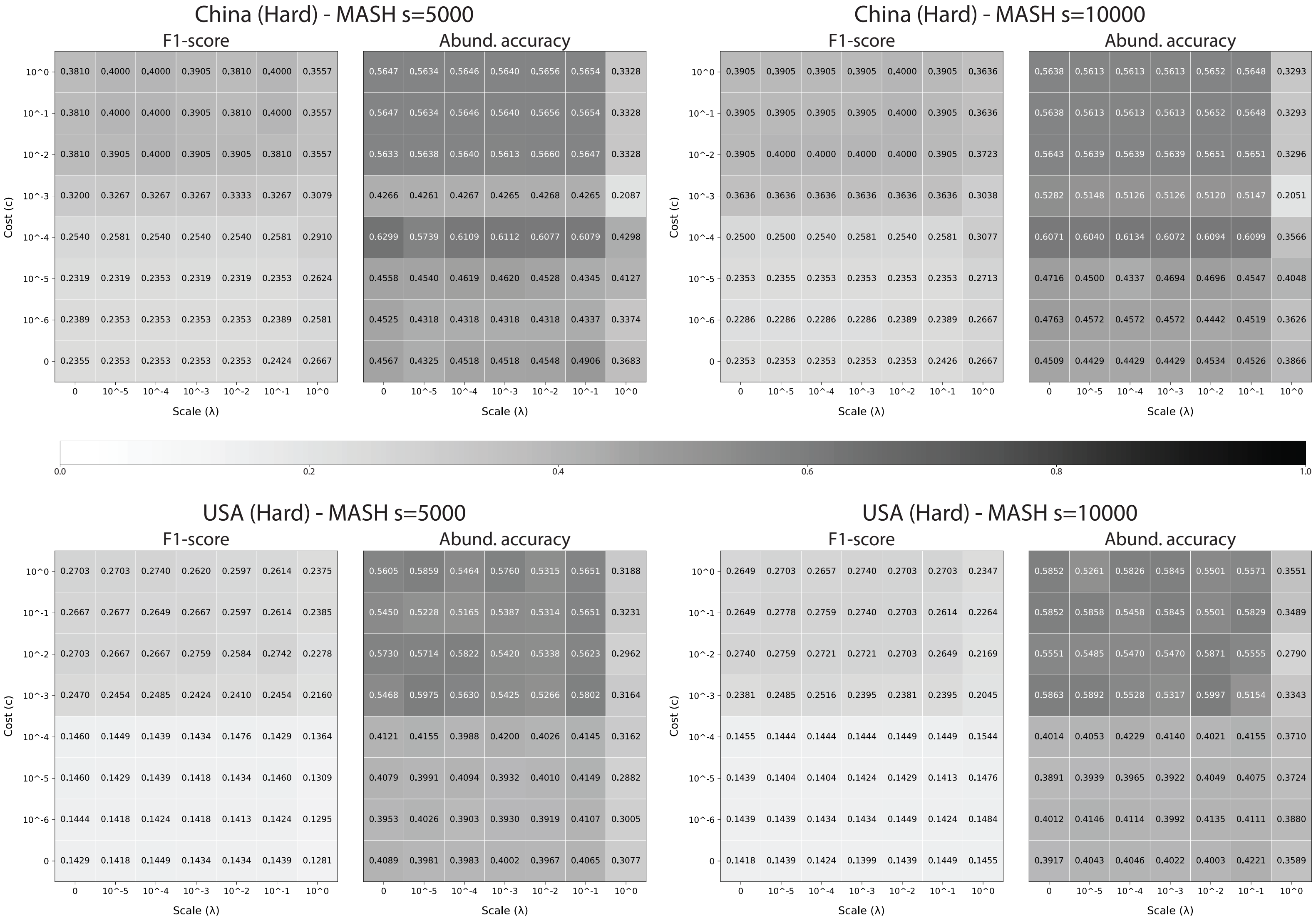

**Supplementary Figure S3:** Accuracy metrics for Hard samples across selection cost $c$ and scale parameter $\lambda$ during SARS-CoV-2 tuning experiments using MASH to estimate distances.

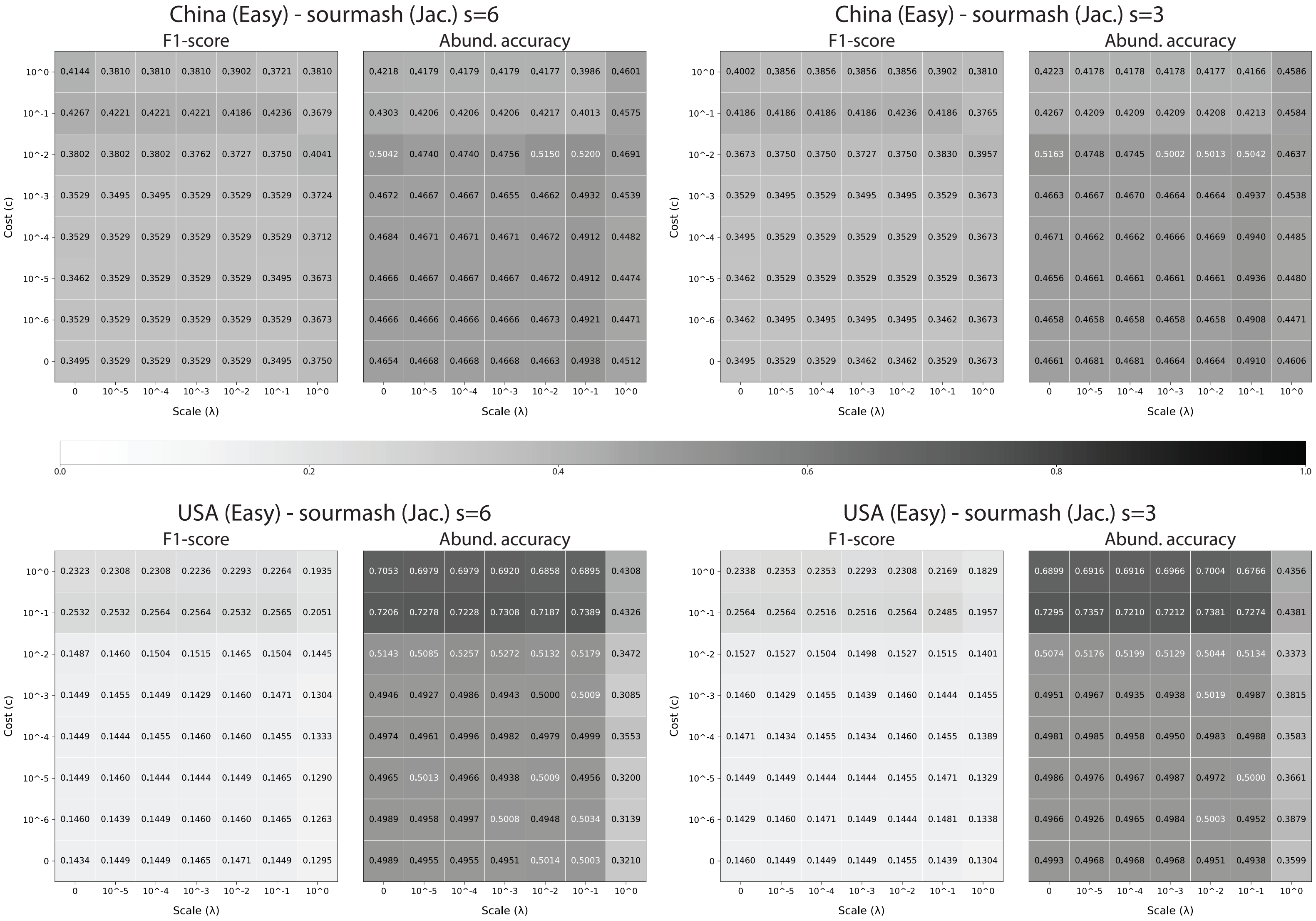

**Supplementary Figure S4:** Accuracy metrics for Easy samples across selection cost $c$ and scale parameter $\lambda$ during SARS-CoV-2 tuning experiments using sourmash (Jaccard) to estimate distances.

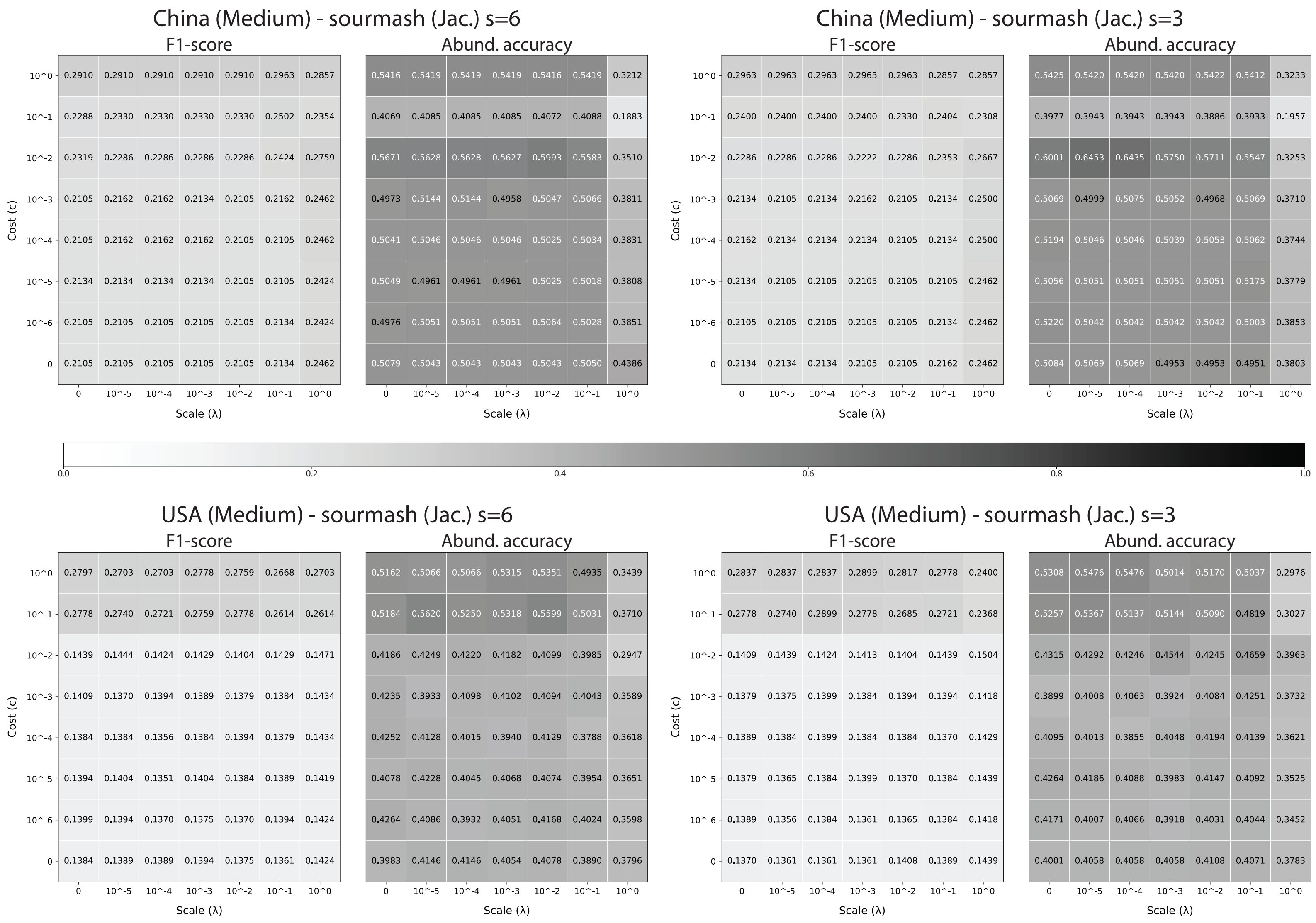

**Supplementary Figure S5:** Accuracy metrics for Medium samples across selection cost $c$ and scale parameter $\lambda$ during SARS-CoV-2 tuning experiments using sourmash (Jaccard) to estimate distances.

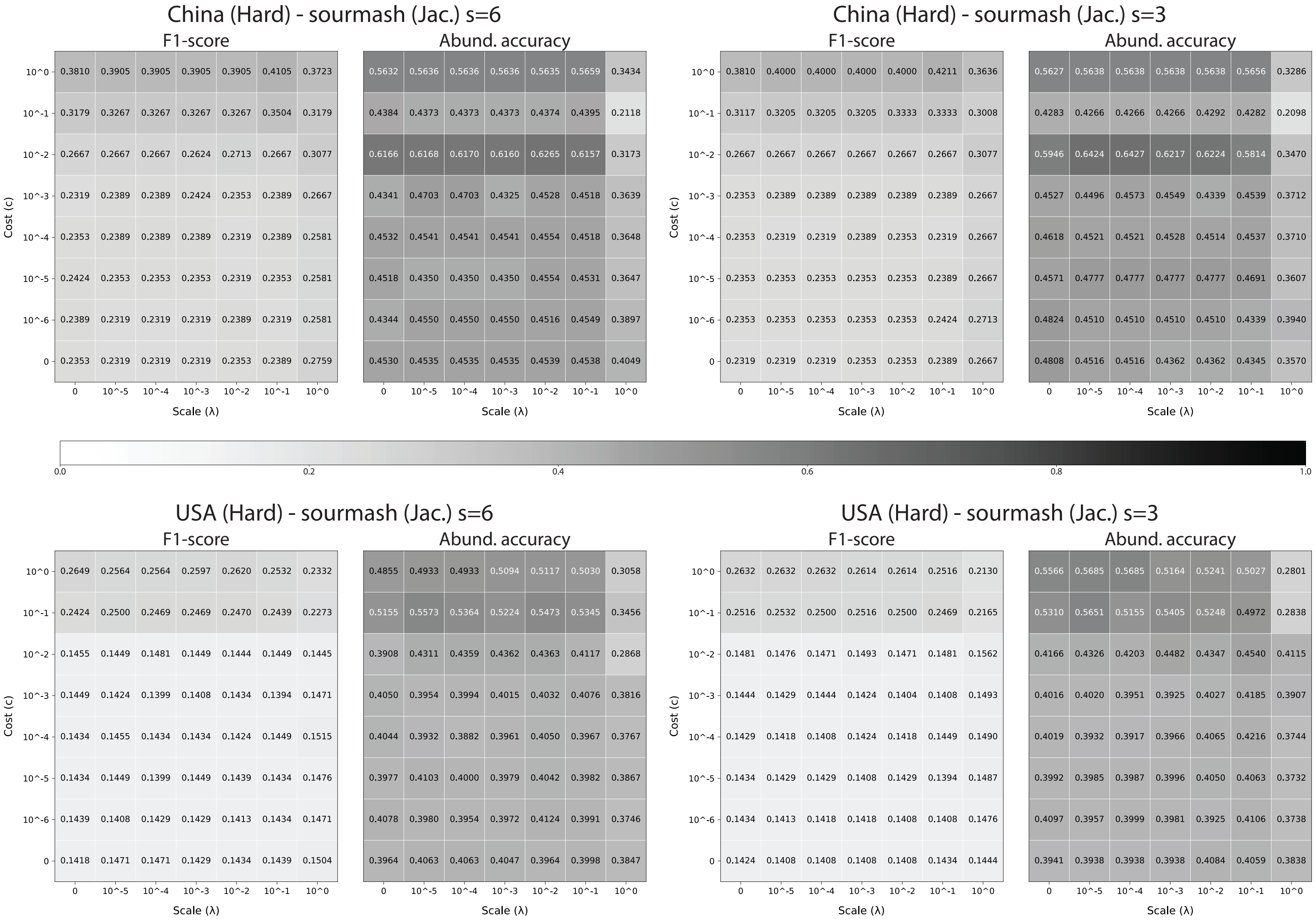

**Supplementary Figure S6:** Accuracy metrics for Hard samples across selection cost $c$ and scale parameter $\lambda$ during SARS-CoV-2 tuning experiments using sourmash (Jaccard) to estimate distances.

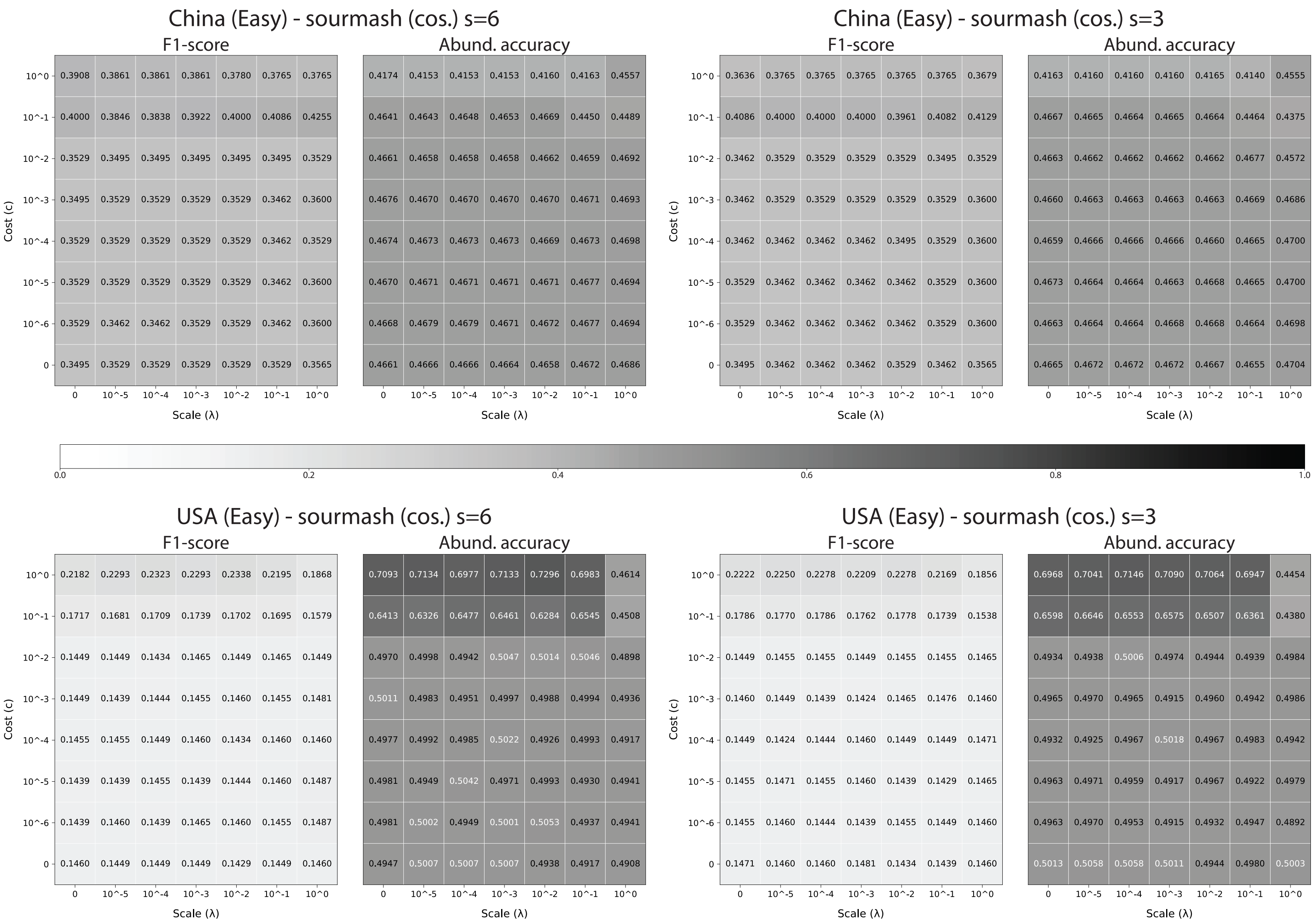

**Supplementary Figure S7:** Accuracy metrics for Easy samples across selection cost $c$ and scale parameter $\lambda$ during SARS-CoV-2 tuning experiments using sourmash (cosine) to estimate distances.

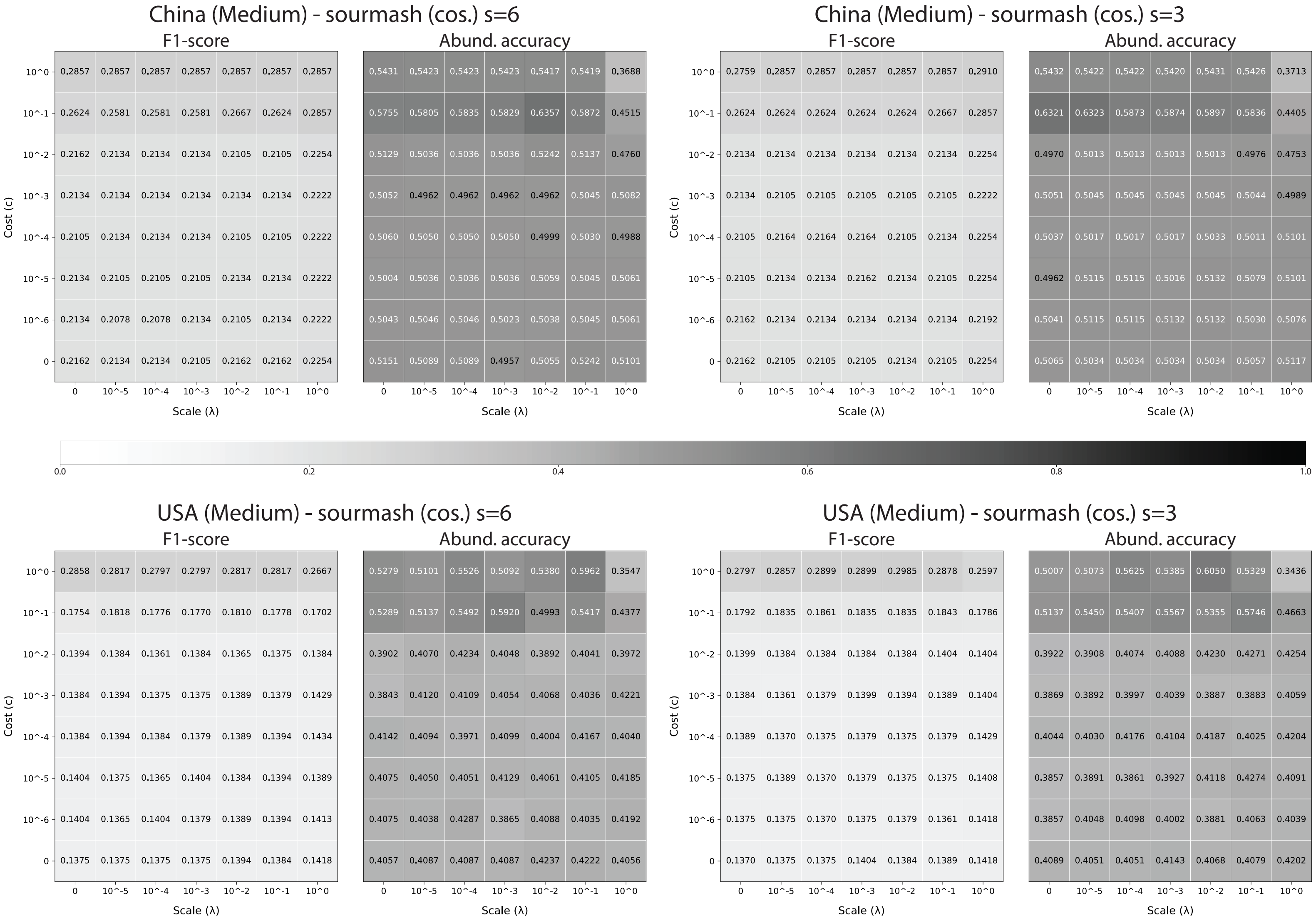

**Supplementary Figure S8:** Accuracy metrics for Medium samples across selection cost $c$ and scale parameter $\lambda$ during SARS-CoV-2 tuning experiments using sourmash (cosine) to estimate distances.

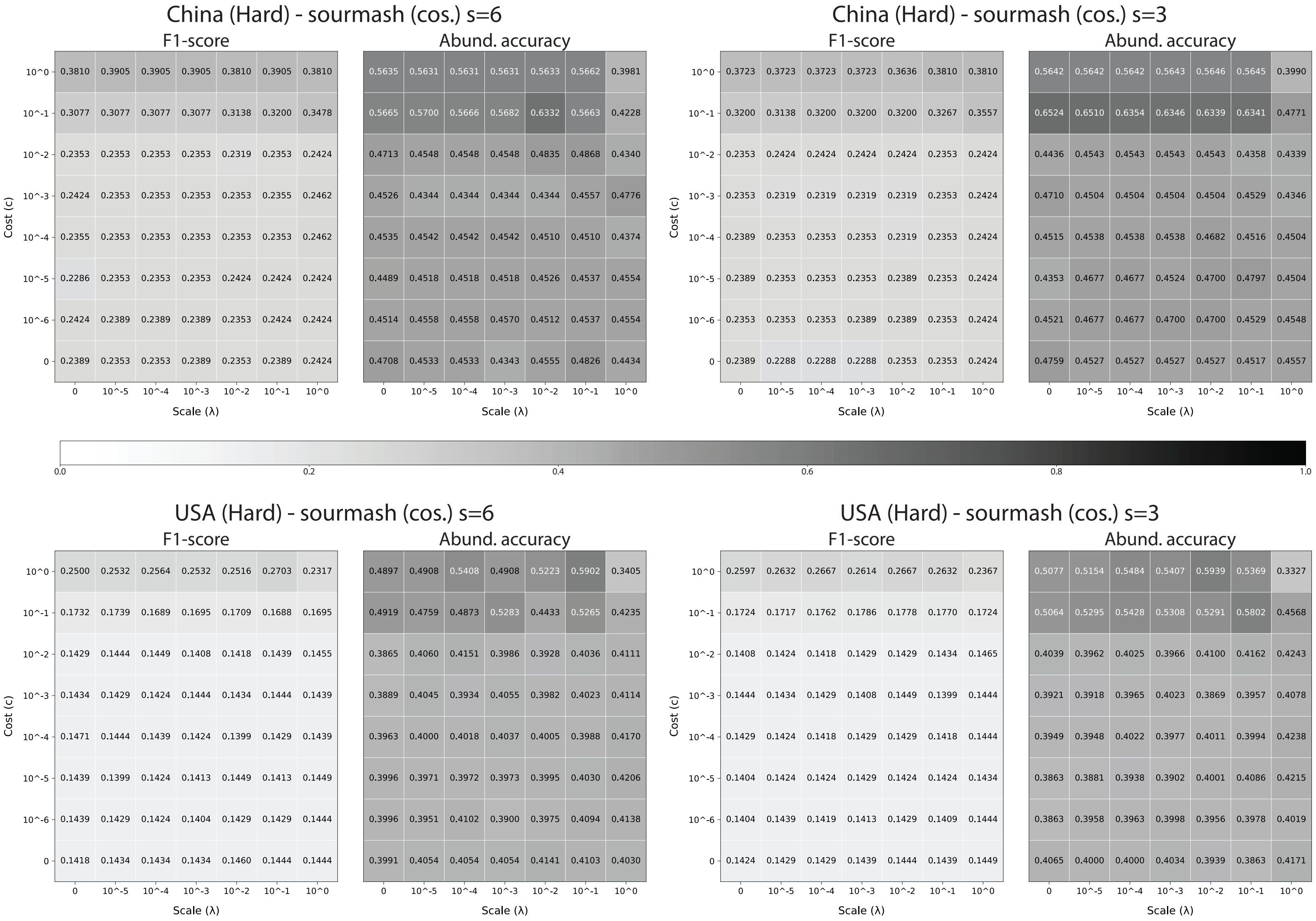

**Supplementary Figure S9:** Accuracy metrics for Hard samples across selection cost $c$ and scale parameter $\lambda$ during SARS-CoV-2 tuning experiments using sourmash (cosine) to estimate distances.
